# Evolution of the complex transcription network controlling biofilm formation in *Candida* species

**DOI:** 10.1101/2020.11.08.373514

**Authors:** Eugenio Mancera, Isabel Nocedal, Stephen Hammel, Megha Gulati, Kaitlin F. Mitchell, David R. Andes, Clarissa J. Nobile, Geraldine Butler, Alexander D. Johnson

**Affiliations:** Departamento de Ingeniería Genética, Centro de Investigación y de Estudios Avanzados del Instituto Politécnico Nacional, Unidad Irapuato, Irapuato, Mexico; Department of Microbiology and Immunology, University of California San Francisco, San Francisco, CA, USA; Department of Biology, Massachusetts Institute of Technology, Cambridge, USA; School of Biomolecular and Biomedical Science, Conway Institute, University College Dublin, Dublin 4, Ireland; The School of Computer Sciences and IT, Western Gateway Building, University College Cork, Ireland; Department of Molecular and Cell Biology, University of California, Merced, CA, USA; Molecular Cell, Cell Press, Cambridge, MA, USA; Department of Medical Microbiology and Immunology, University of Wisconsin, Madison, United States, USA

**Keywords:** transcriptional evolution, biofilm, *Candida albicans*, *Candida dubliniensis*, *Candida tropicalis*, *Candida parapsilosis*

## Abstract

We examine how a complex transcription network composed of seven “master” regulators and hundreds of target genes evolved over a span of approximately 70 million years. The network controls biofilm formation in several *Candida* species, a group of fungi that are present in humans both as constituents of the microbiota and as opportunistic pathogens. The ability to form biofilms is crucial for microbial colonization of different host niches, particularly when an implanted medical device is present. We examined and compared the network underlying biofilm formation across four *Candida* species (*C. albicans, C. dubliniensis, C. tropicalis, and C. parapsilosis*), all of which form biofilms composed of multiple cell types. To describe the salient features of the network across different species, we employed four approaches: (1) we phenotypically characterized the biofilms formed by these species using a variety of methods; (2) we knocked out — one by one — the master regulators identified in *C. albicans* in the four species and monitored their effect on biofilm formation; (3) we identified the target genes of 18 master regulator orthologs across the four species by performing ChIP-seq experiments; and (4) we carried out transcriptional profiling across each species during biofilm formation. Additional network information was obtained by analyzing an interspecies hybrid formed between the two most closely related species, *C. albicans* and *C. dubliniensis*. We observed two major types of changes that have occurred in the biofilm circuit since the four species last shared a common ancestor. Master regulator “substitutions” occurred over relatively long evolutionary times, resulting in different species having overlapping, but different sets of master regulators of biofilm formation. Second, massive changes in the connections between the master regulators and their target genes occurred over much shorter timescales. Both types of change are crucial to account for the structures of the biofilm networks in extant species. We believe this analysis is the first detailed, empirical description of how a complex transcription network has evolved.

## INTRODUCTION

Many of the most medically relevant fungi belong to the *Candida* genus. These microbes are part of the human microbiota, but under specific circumstances — such as imbalances in components of the microbiota or suppression of the immune system of the host — they can proliferate as opportunistic pathogens and cause disease (Calderone and Clancy 2012; Turner and Butler 2014; Kullberg and Arendrup 2015; Romo and Kumamoto 2020). These diseases, which were already documented by the ancient Greeks, range from mild cutaneous disorders to systemic infections with high mortality rates (Lynch 1994; Calderone and Clancy 2012; Kullberg and Arendrup 2015; Nobile and Johnson 2015). Although they are usually studied in planktonic (suspension) cultures in the laboratory, *Candida* species, like many microbes, are often found in nature as biofilms, communities of cells associated with surfaces. For *Candida albicans*, the best studied and most clinically relevant of the *Candida* species, biofilms consist of a lower sheet of cells in the yeast form (spherical, budding cells) overlaid by a layer of filamentous cells (hyphae and pseudohyphae) and surrounded by an extracellular matrix composed of proteins and secreted polysaccharides (Blankenship and Mitchell 2006; Nobile and Johnson 2015; Lohse *et al*. 2018). The matrix, together with specific gene expression changes within biofilms (for example, the upregulation of drug efflux pumps), provide protection from environmental stresses including antifungal drug treatment. The ability of *C. albicans* to form biofilms has been associated both with its versatility in occupying different niches in the human host and its inherent resistance to antifungal drugs. These features are especially important for individuals with implanted medical devises, which provide substrates for biofilm formation and where often the only effective treatment is replacement of the device (Donlan 2001). Biofilms also shed live yeast-form cells and thereby serve as reservoirs for further colonization in the human body (Nobile and Johnson 2015).

*C. albicans* biofilm formation begins with the adhesion of yeast cells to a surface, followed by cell division and morphological differentiation to form an upper layer of filamentous cells. The biofilm matures through the secretion of the extracellular matrix (Blankenship and Mitchell 2006; Nobile and Johnson 2015; Lohse *et al*. 2018). In *C. albicans*, a complex transcription network regulates this process; it consists of seven “master” transcription regulators (Bcr1, Brg1, Efg1, Flo8, Ndt80, Rob1 and Tec1) that control each other’s expression and, collectively, bind to the control regions of more than a thousand target genes — around one sixth of the total number of genes present in the genome of this species (Figure 1) (Nobile *et al*. 2012; Fox *et al*. 2015). Despite the complexity of the biofilm regulatory netwrok, several lines of evidence suggest that this network originated relatively recently. For example, genes that are highly expressed during biofilm formation are enriched for genes that are relatively young, meaning that they only have a clear ortholog in species closely related to *C. albicans* (Nobile *et al*. 2012). Apart from the literature available for *C. albicans*, most of the work to understand biofilm formation in *Candida* species has been carried out with *Candida parapsilosis* (Ding and Butler 2007; Connolly *et al*. 2013; Holland *et al*. 2014). *C. parapsilosis* diverged from a last common ancestor with *C. albicans* nominally 70 million years ago (Mishra *et al*. 2007; Butler *et al*. 2009). Although six of the seven master regulators of biofilm formation in *C. albicans* have clear orthologs in *C. parapsilosis*, only two of them are required for biofilm formation in the latter species (Holland *et al*. 2014). *Candida dubliniensis* and *C. tropicalis* are more closely related to *C. albicans* (see Figure 2) and are also known to form biofilms (Ramage *et al*. 2001; Silva *et al*. 2011; Pujol *et al*. 2015; Araujo *et al*. 2017; Dominguez *et al*. 2018; Kumari *et al*. 2018), but the regulatory circuits that control this process are largely unknown.

**Figure 1.**
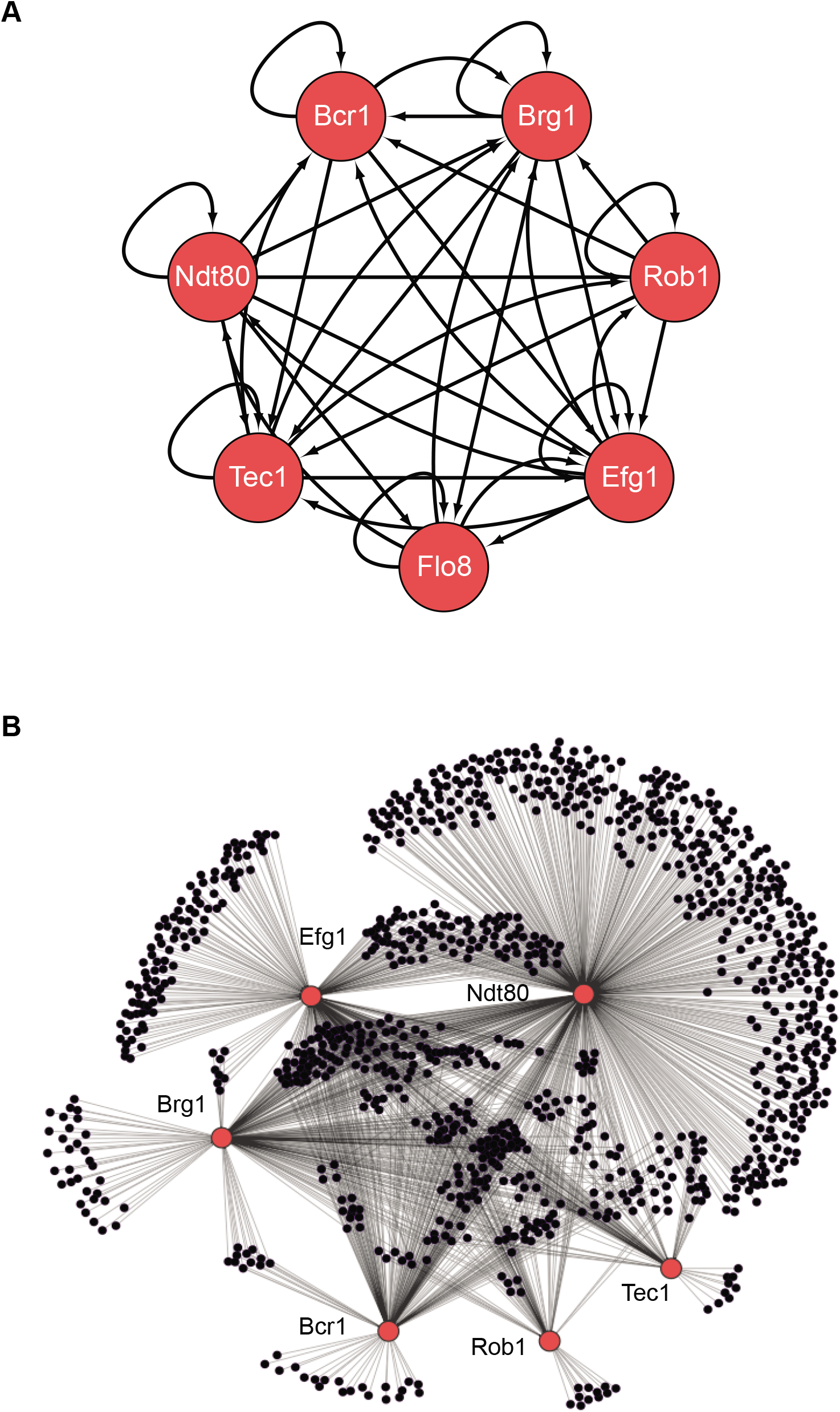
The biofilm transcription network in *Candida albicans*. (A) The seven master transcription regulators identified in genetic screens (Nobile *et al*. 2012; Fox *et al*. 2015), and the interactions among them as determined by ChIP-seq and ChIP-qPCR. (B) Binding interactions (determined by ChIP-seq) between the master regulators (red) and their target genes (black). Many target genes are bound by more than one regulator. Note that genome-wide binding data is not available for Flo8, and thus it is missing from the larger network diagram.

**Figure 2.**
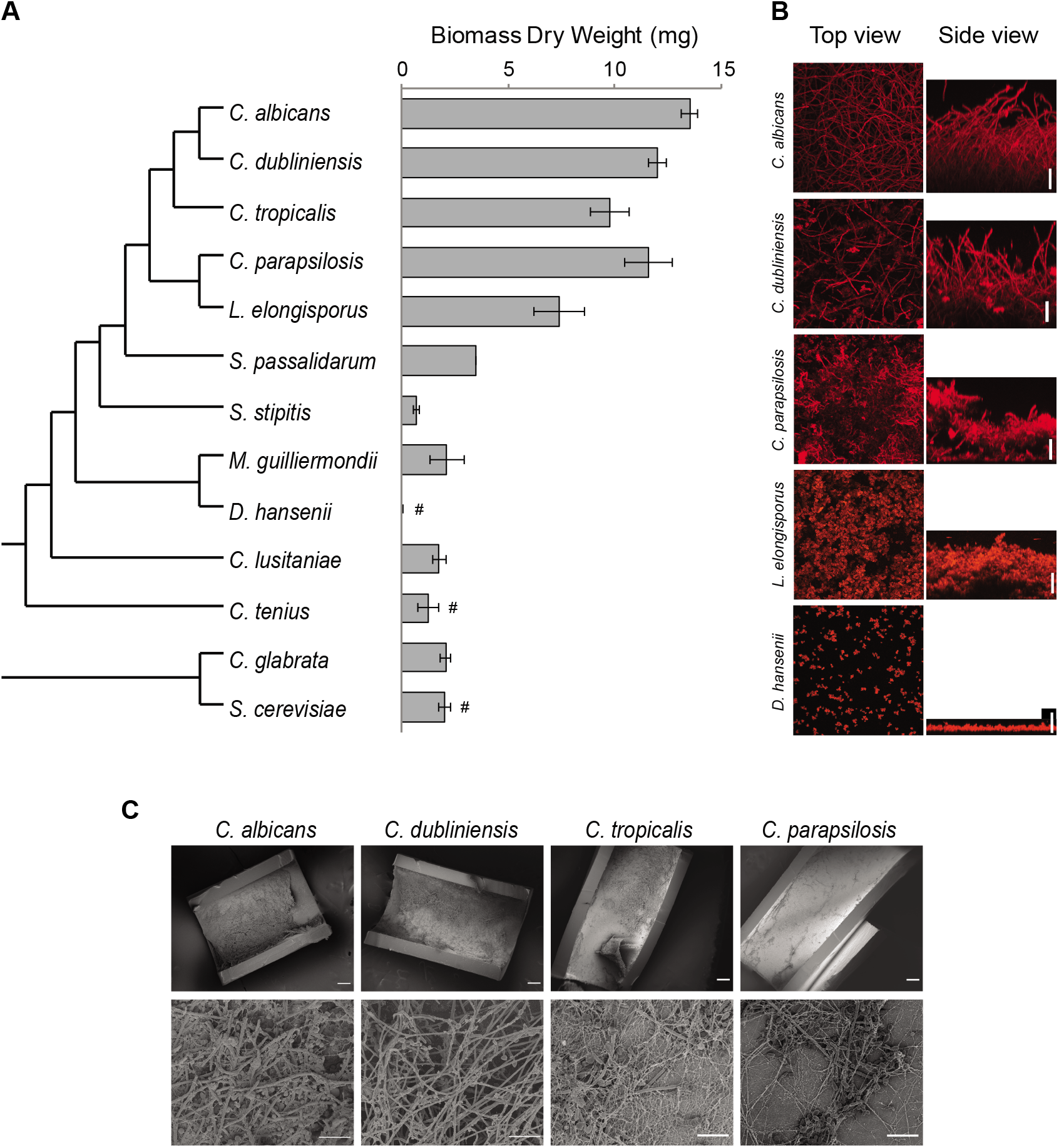
Diversity in biofilm formation across fungal species. (A) Biofilm biomass dry weight was determined for different fungal species grown on the bottoms of a polystyrene 6-well plates in Spider 1% glucose medium at 37°C for 48 hrs. The mean and standard deviation was calculated from five replicates. Hashtags denote species that do not grow well at 37°C and for which biofilms were grown at 30°C. The cladogram to the left shows the phylogenetic relationship of the species (Byrne and Wolfe 2005; Maguire *et al*. 2013). All species analyzed belong to the CTG-Ser1 clade apart from *C. glabrata* and *S. cerevisiae*. (B) Morphology of biofilms formed by five representative CTG clade species visualized by CSLM. Biofilms were grown as described above, but on the surface of silicone squares. Scale bars represent 50 μm. (C) Biofilm formation by *Candida* species in an *in vivo* rat catheter model (Andes *et al*. 2004). Biofilms were grown for 24 hrs. and were visualized by scanning electron microscopy (SEM). Two magnifications are shown in the lower and upper panel for each species and the scale bars represent 100 and 20 μm, respectively.

To understand how the complex transcription network that controls biofilm formation evolved, we began with the seven master regulators of biofilm formation in *C. albicans* and determined whether their orthologs also controlled biofilm formation in *C. dubliniensis* and *C. tropicalis*. Using ChIP-seq, we mapped the targets of the orthologs in *C. dubliniensis, C. tropicalis* and *C. parapsilosis*. A comparison of the extant networks showed that the two main components of the network, master regulators and target genes, moved in and out of the network at very different rates over evolutionary time. While the regulators moved gradually and in rough correlation with small phenotypic changes in biofilm structure, the master regulator-target gene connections changed very quickly. The large-scale changes in connections observed between closely related species did not appear to have a major impact on biofilm phenotypes, at least as monitored *in vitro*. These results suggest an evolutionary route through which complex regulatory networks could rapidly explore new network configurations (and perhaps new phenotypes) without disrupting existing functions.

## RESULTS

### Only closely related species to *C. albicans* form complex biofilms

To understand how the transcription network that controls biofilm formation changed over evolutionary timescale, we first phenotypically characterized the biofilms formed *in vitro* by the different species of the so-called CTG clade. This clade, which includes but extends beyond *Candida* species, was traditionally named CTG due to its unusual property of decoding the CTG codon as serine instead of the usual leucine (Figure 2). Recently, this clade has been renamed CTG-Ser1 because other Ascomycota clades were discovered to also have unusual codon usage (Krassowski *et al*. 2018). To define an optimal growth medium for these assays we tested biofilm formation under several conditions typically used to study biofilms of *Candida* species (Garcia-Sanchez *et al*. 2004; Richard *et al*. 2005; Kucharikova *et al*. 2011; Nobile *et al*. 2012; Lohse *et al*. 2017). For the initial tests, we focused on *C. albicans* and the three species that are most closely related to it and that commonly inhabit humans, *C. dubliniensis, C. tropicalis* and *C. parapsilosis* (Figure 2) (Turner and Butler 2014; Gabaldon *et al*. 2016). The estimated divergence time for *C. dubliniensis, C. tropicalis* and *C. parapsilosis* from the last common ancestor with *C. albicans* is approximately 20, 45, and 70 million years, respectively (Mishra *et al*. 2007; Butler *et al*. 2009; Moran *et al*. 2012). *C. albicans, C. tropicalis* and *C. parapsilosis* have been previously shown to form biofilms, while less is known about biofilm formation in *C. dubliniensis* (Silva *et al*. 2011; Araujo *et al*. 2017; Dominguez *et al*. 2018; Kumari *et al*. 2018). Biofilms were grown *in vitro* on silicone squares at 37°C for 48 hours with shaking and were monitored by confocal scanning laser microscopy (CSLM), as has been previously described (Nobile *et al*. 2012). We tested eight different growth media, and only Spider medium with glucose (rather than mannitol) as the carbon source allowed all four species to form thick, well-structured biofilms (Sup Table 1). Our results also showed that environmental conditions are important determinants of biofilm formation for some of these species. While *C. albicans* formed thick biofilms in all media tested, *C. tropicalis* biofilm formation, for example, depended very much on carbon source.

Given that there could be differences in the speed at which different species form biofilms, we also assessed biofilm formation as a function of time for the same four species. Biofilms were formed as described above and were monitored at seven different time points from 30 minutes to 96 hours after cell adhesion under the confocal microscope. Although *C. albicans* formed biofilms more rapidly, by 48 hours all four species had formed mature biofilms that did not significantly change at later time points (Sup Figure 1).

Once we had defined an optimal biofilm growth medium (Spider + glucose) and time point (48 hours) we extended our analysis to other species of the CTG clade (Maguire *et al*. 2013). In addition to CSLM as described above, we monitored biofilm formation in two additional ways: we determined the biomass dry-weight of biofilms formed on the bottom of polystyrene plates, and, using a microfluidic flow cell, we continuously monitored biofilm formation by time lapse photography using an optical microscope (Nobile *et al*. 2012; Lohse *et al*. 2017). The three methods are complementary: biomass determination is a quantitative method that reduces biofilm formation to a single number, confocal microscopy is qualitative, but allows detailed characterization of the structure of the biofilm, and the microfluidic assay reveals biofilm formation in real-time under a defined flow; the flow rate was adjusted to mimic that of an average catheter implanted in a vein (Gulati *et al*. 2017; Lohse *et al*. 2017). Because some of the species we tested are known to grow poorly at 37°C, we performed the assays at 30°C for those species (Kurtzman *et al*. 2011). Of the 16 species tested, those closest to *C. albicans* formed the thickest biofilms and, in general, the greater the phylogenetic distance from *C. albicans* the thinner the biofilm formed. Only the two species closest to *C. albicans (C. dubliniensis* and *C. tropicalis*) formed biofilms that are structurally very similar to *C. albicans* biofilms, with a basal layer of yeast cells underlying a thick layer of filamentous cells (hyphae and pseudohyphae) enclosed in an extracellular matrix. Biofilms formed by *C. parapsilosis* appeared similar at low resolution, but a closer examination showed that the layer of filamentous cells is composed largely of pseudohyphae rather than a mixture of true hyphae and pseudohyphae. Under these conditions, *Lodderomyces elongisporus* formed thinner biofilms composed only of yeast cells. Moving further away from *C. albicans, Spathaspora passalidarum, Meyerozyma guilliermondii* and *Clavispora lusitaniae* form even thinner biofilms, while *Scheffersomyces stipites, Debaryomyces hansenii* and *C. tenuis*, did not form biofilms under the conditions we tested; only a few cells were observed adhering to the surface. We also performed CSLM assays in an additional medium (RPMI), and the results generally agreed with those described above. Our results with the microfluidic assays showed a similar trend: only those species that are phylogenetically closest to *C. albicans* were able to rapidly form biofilms under flow conditions in the microfluidic device (Sup Figure 2).

As a reference, we also characterized biofilm formation in two other ascomycetous yeast species that lie outside the CTG clade, *C. glabrata* and *S. cerevisiae*. Although not closely related to the CTG clade (despite its name), *C. glabrata* is an important opportunistic human pathogen, while *S. cerevisiae* is used extensively in the food and beverage industries and is a widely employed model organism. As can be seen in Figure 2, neither of these species formed biofilms that resembled those formed by *C. albicans* and its close relatives in the assays and conditions that we tested.

To assess whether the results observed *in vitro* can be recapitulated *in vivo*, we tested the ability of *C. albicans, C. dubliniensis, C. tropicalis* and *C. parapsilosis* to form biofilms in the rat catheter model, a well-established *in vivo* biofilm model (Andes *et al*. 2004). All four species were able to form biofilms, although the biofilms formed by *C. albicans* and *C. dubliniensis* were considerably thicker and more filamentous (Figure 2C). These results agree with previous *in vivo* characterizations performed for *C. albicans* and *C. parapsilosis* (Nobile *et al*. 2012; Connolly *et al*. 2013) and provide new information on the intermediate species.

In summary, our results show that the ability to form biofilms that resemble those of *C. albicans* is limited to its most closely related species. In terms of biomass, there is a sharp drop off outside *C. parapsilosis* while, in terms of biofilm structure, only *C. dubliniensis* and *C. tropicalis* form biofilms similar to those of *C. albicans*, in terms of all three morphological cell types being represented. Of all the species studied, *C. albicans* biofilm formation is the most rapid and most robust to environmental changes; moreover, the biofilms formed by this species are the most stable to physical manipulation (results not shown).

### The regulatory core of the biofilm transcription network changed gradually over time

To gain insight into the evolutionary changes that occurred in the transcription network that controls biofilm formation at a molecular level, we first studied the function of the seven master regulators of the *C. albicans* network (Figure 1). Given the phenotypic results described above, we centered the analysis on *C. albicans, C. dubliniensis, C. tropicalis* and *C. parapsilosis*. All four of these species are common in humans (Turner and Butler 2014), and the first three form similar structural types of biofilms. As described above, *C. parapsilosis* also forms biofilms, but its biofilms show more pronounced differences. All seven master regulators of the network in *C. albicans* (Figure 1) have clear orthologs in the other three closely related species, with the exception of Rob1. Rob1 has a patchy phylogenetic distribution, with syntenic orthologs present in *C. albicans, C. dubiniensis* and *C. tropicalis*, but apparently absent from *C. parapsilosis* and closely related species. However, Rob1 orthologs are present in other more distantly related CTG species, which supports the hypothesis that Rob1 was either lost or was evolving sufficiently rapidly in the *C. parapsilosis* linage that it cannot be recognized (Maguire *et al*. 2013).

To test whether the orthologs of the *C. albicans* master regulators are involved in biofilm formation in the other species, we generated gene deletion knockouts in *C. dubliniensis* and *C. tropicalis*. The knockouts in *C. albicans* and *C. parapsilosis* had been previously generated as part of large transcription regulator deletion projects (Homann *et al*. 2009; Holland *et al*. 2014). To make the knockouts in *C. dubliniensis* and *C. tropicalis* we used amino acid auxotrophic strains and employed a gene knockout strategy similar to the that previously used for *C. albicans* and *C. parapsilosis* (Mancera *et al*. 2019).

The ability of the different gene knockout strains to form biofilms was monitored by biomass dry-weight determination and by CSLM. As can be observed in Figure 3 and Sup Fig. 3, all seven master regulators identified in *C. albicans* were also required for biofilm formation in *C. dubliniensis*. The results were different for *C. tropicalis*; here, only five of the seven were required, with Rob1 and Flo8 appearing dispensable for biofilm formation under our laboratory conditions. The biofilms formed by the *ROB1* and *FLO8* deletion mutants in *C. tropicalis* were actually slightly heavier and the hyphal layer was denser than biofilms formed by the parental (wildtype) strain, suggesting that these two regulators may have assumed a subtle inhibitory role in these species. For *C. parapsilosis*, it was previously shown that, of the seven master biofilm regulators in *C. albicans*, only Bcr1 and Efg1 are indispensable for biofilm formation. The Ntd80 deletion in *C. parapsilosis* fails to form biofilms but also has a general growth defect, making it difficult to ascertain its precise role. Previous work also established that deletion of *CZF1, UME6, CPH2, GZF3* and *ACE2* all exhibit biofilm specific defects in *C. parapsilosis* but not in *C. albicans* (Holland *et al*. 2014). Overall, our results, together with previous observations, show that the group of master regulators underlying biofilm formation in *C. albicans* has diversified in other *Candida* species with the degree of diversity roughly paralleling their evolutionary distance from *C. albicans*.

**Figure 3.**
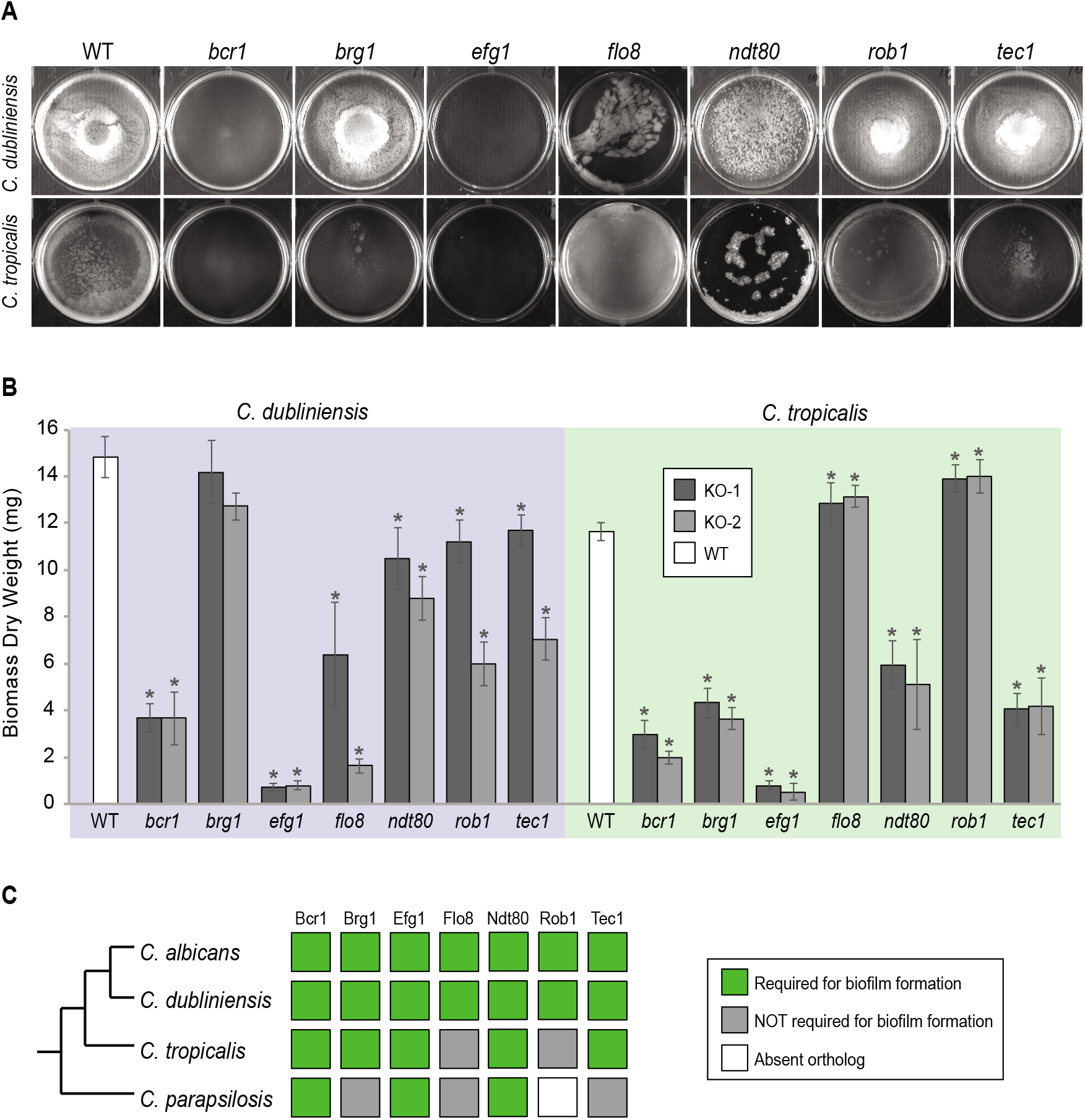
Roles of orthologs of the seven *C. albicans* master regulators in biofilm formation. (A) Phenotypic characterization of biofilms formed by the gene deletion knockouts of orthologs of the seven master *C. albicans* biofilm regulators. Images show biofilms grown on the bottoms of a polystyrene 6-well plate in Spider 1% glucose medium at 37°C for 48 hrs. (B) Dry weights of biofilms formed by the gene deletion mutants grown as described in (A). The means and standard deviations were calculated from five replicates for two independent gene deletion knockout isolates (KO-1 and KO-2). Asterisks denote statistically significant different weights when compared to the corresponding parental strain using a Student’s two-tailed paired *t* test (*P* < 0.05). (C) Summary diagram showing the conservation of the seven master regulators in biofilm formation across the three most closely related species to *C. albicans*. The data for *C. albicans* was obtained from (Nobile *et al*. 2012; Fox *et al*. 2015), and that for *C. parapsilosis* from (Holland *et al*. 2014).

### Target genes of the master biofilm regulators differ greatly among *Candida* species

To determine how the binding connections between the biofilm master regulators and their target genes have changed over evolutionary time, we determined genome-wide protein-DNA interactions of the master regulators in *C. albicans, C. dubliniensis, C. tropicalis* and *C. parapsilosis* by chromatin immunoprecipitation followed by next-generation sequencing (ChIP-seq) (Johnson *et al*. 2007). To this end we tagged each of the regulators with a Myc epitope tag that can be immunoprecipitated using a commercially available antibody. This strategy has the advantage that a single antibody with the same affinity could be used in all experiments, and control experiments could be performed with untagged strains. All the ChIP-seq experiments were performed in mature biofilms grown for 48 hours at 37°C in the optimal medium described above. For unknown technical reasons, not all regulators could be immunoprecipitated in all species. Of all the ChIP-seq experiments performed, protein-DNA interactions could be reliably determined for 18 regulators across the four species (Sup Table 2). We believe that this coverage, although not complete, is more than sufficient to uncover the general trends of biofilm network evolution.

The large amount of new genome-wide protein-DNA interaction data reported here will be useful in future studies of biofilm formation in these *Candida* species. Very few “structural” genes have been implicated in biofilm formation, and the ChIP-seq results across species could greatly facilitate the identification of key non-regulatory genes required for biofilm formation. For example, specific master regulator-target gene connections that are preserved across multiple species may point to target genes that are especially important for biofilm formation. Such a hypothesis could be tested by knocking out the target gene of interest. In the following sections, we consider, in general terms, how the biofilm network (defined as the set of master regulators and connections between them and their target genes) differs among *Candida* species and what these differences reveal about the evolution of the network.

To compare gene targets between species, we assigned each ChIP occupancy site to the ORF with the nearest downstream start codon. To compare the changes in master regulator-target gene connections across species, were first examined the target genes that are conserved across the species. The overall percentage of one-to-one orthologs between the four species ranged from a high of 91% between *C. albicans* and *C. dubliniensis* to a low of 74% between *C. tropicalis* and *C. parapsilosis* (Maguire *et al*. 2013). If we consider only the one-to-one orthologs, it becomes clear that the connections between master regulators and conserved target genes vary greatly across these species (Figure 4). Between the two most closely related species (*C. albicans* and *C. dubliniensis*), the master regulator that showed the highest conservation of target gene connections was Ndt80, but the overlap was only about 50%. Between *C. albicans* and *C. parapsilosis*, this value drops to about 25%. Although only 12% of Ndt80 target gene connections are common to all four species, the overlap for each species pair is larger than expected by chance (hypergeometric test, *P* < 0.05), indicating that a small but significant group of Ntd80-target gene connections have been preserved across these species. The other master regulators show an even lower degree of conserved regulator-target gene connections. For example, Rob1 shares only about 12% of its target gene connections between the two most related species, *C. albicans* and *C. dubliniensis*.

**Figure 4.**
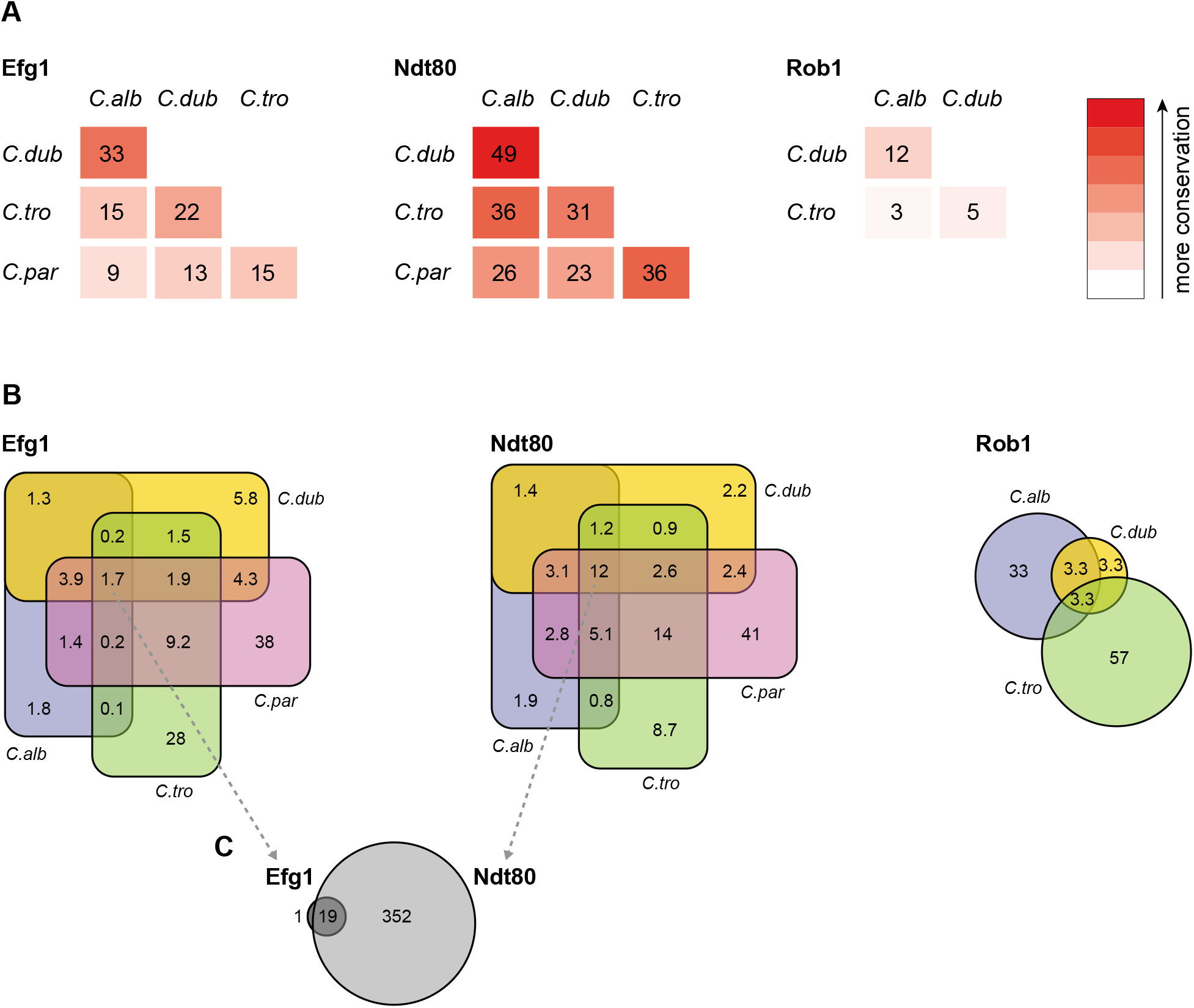
Connections between master regulators and target genes are highly divergent across species. (A) Pair-wise comparison of shared target genes for Ndt80, Efg1 and Rob1 between species. Target genes were determined by ChIP-seq as detailed in the methods. The numbers represent the percentage of overall target genes conserved between each pair of species considering only genes that have orthologs in the two species. Note that Rob1 is absent in *C. parapsilosis* (Maguire *et al*. 2013). (B) Venn diagrams depicting the overlap of regulator-target gene connections across species, considering only genes that have orthologs in all four species for Efg1 and Ndt80 and considering genes that have orthologs in *C. albicans, C. dubliniensi* and *C. tropicalis* for Rob1. Numbers in each section of the diagrams represent the percentage of master regulator-target gene connections, with the total number of connections for each regulator set at 100%. Note that for Efg1 and Ndt80 the size of the color sections do not correspond to the percentage. (C) Venn diagram depicting the overlap between target genes of Efg1 and Ndt80, considering only target gene orthologs that are present in all four species. Numbers represent the gross number of genes. The diagram indicates that, for genes that are targets of Efg1 and Ndt80 in all species, most Efg1-target gene connections are also Ndt80-target gene connections, even though the target genes themselves are different across species.

Our results also show that each species has target genes in its biofilm network that lack orthologs in the other three species. In *C. albicans*, 26% of the genes bound by at least one of the biofilm master regulators do not have orthologs in the other three species, compared with 19% for the genome as a whole. This analysis extends the previous observation that genes that are overexpressed during biofilm formation in *C. albicans* are often “young” genes, that is genes that lack orthologs in related species (Nobile *et al*. 2012). We observed a similar enrichment of unique genes in the biofilm network of *C. parapsilosis*, but not for *C. dubliniensis* and *C. tropicalis;* in the latter two species the fraction of non-orthologous genes in the biofilm network was approximately the same as that observed for the whole genome. These observations indicate that the biofilm networks of *C. albicans* and *C. parapsilosis* have been more dynamic in recent evolutionary time than those of the other two species.

### High connectivity of the biofilm network is observed in all four species

The initial characterization of the biofilm transcription circuit in *C. albicans* showed that many target genes were directly connected (by binding) to more than one master regulator (Nobile *et al*. 2012), and this general feature of the network architecture is observed across the *Candida* species studied here, despite the low conservation of individual regulator-target gene connections (Figure 4). Perhaps the most notable example is seen by comparing the target genes of Ndt80 and Efg1 across the four species. The set of target genes of Ndt80 is considerably larger, but over 75% of the Efg1 target genes are also Ndt80 targets in all four species. In addition, the binding motifs of these two regulators are enriched in each other’s binding locations in all four species studied (Figure 5A). These observations indicate that, in all species, Efg1 binds in conjunction with Ndt80 even though the target genes of the regulator combination differ greatly across species. The association between Efg1 and Ndt80 agrees with previous planktonic ChIP-seq experiments of Efg1 performed in *C. parapsilosis* where the most enriched binding motif found was that of Ndt80 (Connolly *et al*. 2013). Taken together, these results suggest that the Efg1-Ndt80 association is ancient with respect to the *Candida* species studied here and that it remains preserved across them despite large species to species differences in the target genes bound by the two regulators. Analysis of the other master regulators indicates that combinational control of target genes is very common in all species, although the other examples do not seem as deeply conserved as the Efg1-Ndt8o example.

**Figure 5.**
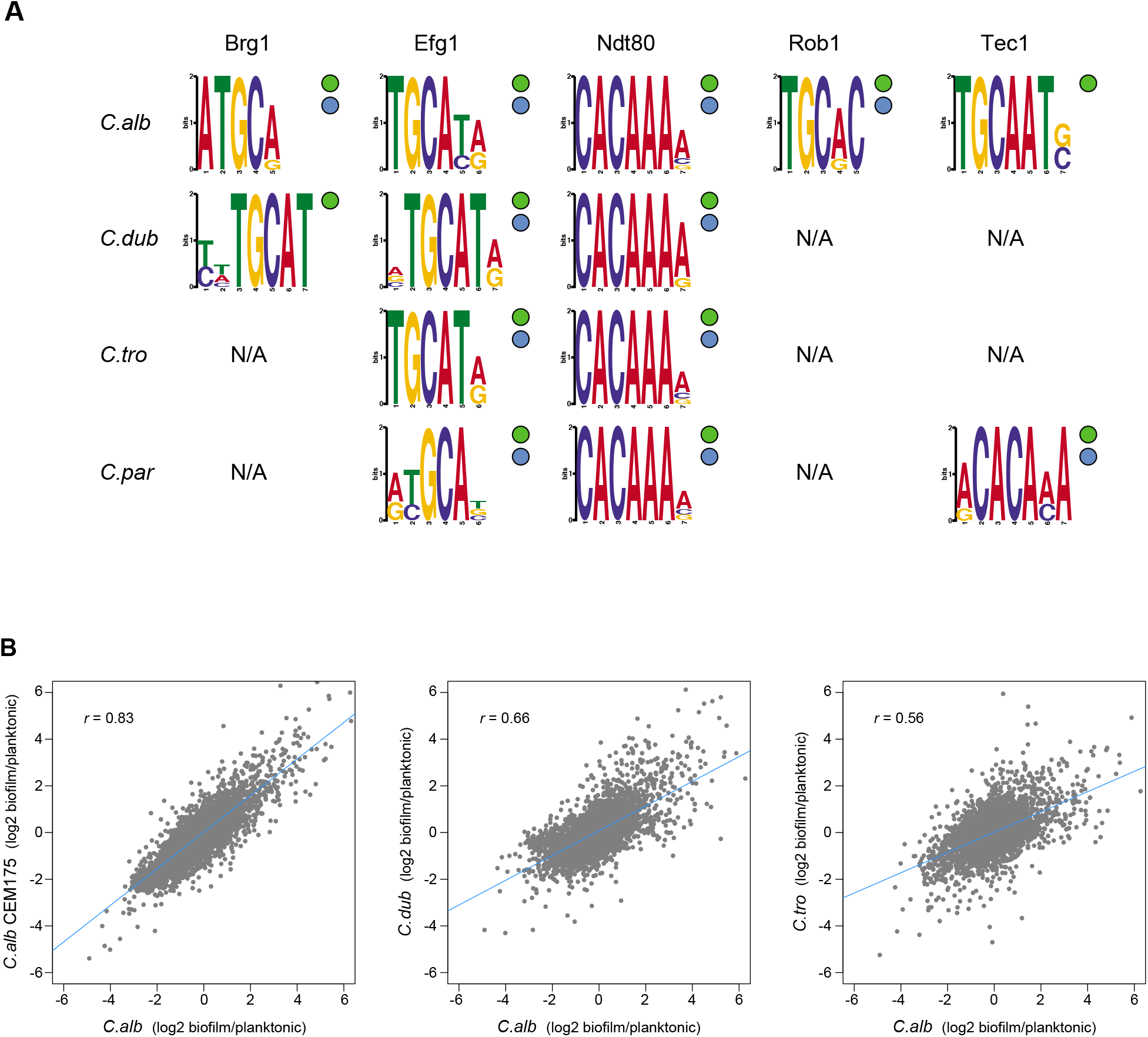
Master regulators retain their DNA-binding specificity while there is considerable variation in gene expression across species. (A) Logos of the most enriched motif in the binding locations of the different master regulators, determined by ChIP-seq across species. The two circles to the right of each logo show whether the Efg1 (green circle) or Ndt80 (blue circle) previously known motifs are enriched in each set of regulator binding locations. (B) Pairwise comparison of transcription profiles under biofilm forming conditions (a time point of 48 hours) of *C. albicans* against *C. dubliniensis* and *C. tropicalis*. As a reference, the comparison between two isolates of *C. albicans* is shown in the left panel. Biofilm specific expression changes were calculated comparing gene expression between biofilm growing conditions and planktonic cultures in the same media. Linear regressions are shown in blue for each comparison.

In terms of overall network structure across species, we also examined whether, as is the case for *C. albicans*, the master regulators bind to their own control region as well as those of the other master regulators. The extent of conservation of the binding connections between one master regulator and the others varies from regulator to regulator (Sup Table 4), with Ndt80 showing the highest conservation. In all four species, Ndt80 binds to its own control region as well as those of all six other regulators, with the exception of the control region of *FLO8* in *C. albicans* and *ROB1* in *C. dubliniensis*. Efg1, Brg1 and Tec1, in that order, follow Ndt80 in their degree of connection conservation, with Rob1 and Bcr1 having the least conserved set of connections. However, we note that the binding data for these two regulators is also the least complete. Ndt80 and Efg1 each bind to their own control regions in all the four species analyzed, while Brg1 exhibits this interaction only in *C. albicans* and *C. dubliniensis*. The binding of the other regulators to their own upstream intergenic regions appears less conserved. Although the ChIP data is unlikely to capture all relevant interactions, we can analyze the existing data to determine whether the connections between regulators (including a regulator binding to its own control region) are more prevalent than that expected by chance. We randomly choose 2,000 groups of seven genes, the number of master regulators in the network, from the binding sets in each species and counted the total number of interactions with the master regulators. In all four species, the number of connections observed between the regulators is greater than what would be expected by chance (*P* << 0.05). Overall, our findings show that high connectivity between the master regulators is conserved in the four *Candida* species analyzed. Thus, despite the extensive changes in the network across species, the high connectivity among regulators remains a structural feature of the network in each species.

### Differences in master regulator-target gene connections are robust to different methods of comparison

We considered the possibility that our results could be skewed by false positive signals intrinsic to ChIP-seq experiments, even when proper controls are used (Chen *et al*. 2012). To deal with this potential problem, we employed two additional criteria for increasing stringency in our analysis of transcription networks (Nocedal *et al*. 2017). First, we filtered the regions identified as enriched in ChIP signal to include only those regions that also had a high scoring regulator binding motif in the intergenic region. As discussed below, the binding motif for some of the master regulators does not vary significantly across the species we studied. Second, we also incorporated gene expression data to further filter the gene targets to those that, (1) have ChIP enrichment, (2) show the presence of a regulator binding motif in the intergenic region, and (3) whose expression changes under biofilm forming conditions. Although the gross number of regulator-target gene connections decreased as the stringency of the filtering criteria increased, the high proportion of differences across *Candida* species described above did not significantly change (Sup Figure 4).

Another potential caveat which could confound our analysis concerns our conclusions being based on the specific conditions under which we induced biofilm formation. To test whether this concern is significant, we performed the ChIP-seq experiments for Ndt80 in *C. albicans, C. dubliniensis* and *C. tropicalis* in biofilms grown in an alternative growth media. Although, in all species, the regulator-target gene connections differed between the two media (Sup Figure 5), the magnitude of the differences in regulator-target gene connections across species was similar to that of our original analysis.

### Changes in master regulator-target gene interactions are due to changes in *cis*-regulatory regions

A possible mechanistic explanation for the high rates of change among the target genes of the biofilm network is that these changes are due to modifications in the *trans* components of the network, for example, changes in the DNA-binding specificity of a master regulator. This type of change has the potential to dramatically change the network over relatively short evolutionary time scales. To explore this possibility, we examined the ChIP-seq binding data for motifs recognized by the master regulators. Performing *de novo* motif searches, we found that the enriched DNA-binding motifs found in the binding regions of the biofilm master regulators were very similar across the four species we examined (Figure 5). Moreover, the *de novo* generated DNA-binding motifs for Efg1 and Ndt80 are similar to those previously reported for their orthologs in other species (Nobile *et al*. 2012; Nocedal *et al*. 2017). Overall, this analysis shows that the DNA-binding specificity of at least some of the master regulators has not changed significantly across these species and cannot explain the diversity of master regulator-target gene connections observed across the species. This conclusion is further supported by analysis of a *C. albicans-C. dubliniensis* hybrid, as described below.

### Biofilm specific gene expression changed rapidly over evolutionary time

A strong prediction of the vast number of species-to-species differences in master regulator-target gene connections documented above is that the genes transcriptionally induced during biofilm formation should differ substantially among species. To test this prediction, we generated genome-wide transcription profiles of *C. albicans, C. dubliniensis* and *C. tropicalis* under biofilm forming conditions. To reveal biofilm-specific changes, we compared these profiles to expression data obtained when the species were grown in suspension cultures in the same medium. As a reference, we also performed the biofilm expression profile in a second *C. albicans* isolate. As seen in Figure 5B, the pairwise differences in the transcription profiles across the three species are significant and reflect their phylogenetic position: the further apart the two species are from one another, the less correlated their transcription profiles are. If we use a lax cut-off of two-fold over-/under-expression to define genes that change their expression during biofilm formation, the overlap between pairs of species is relatively low. For example, only 29% of the genes that change their expression during biofilm formation are shared between *C. albicans* and *C. dubliniensis*, and only 24% are shared between *C. albicans* and *C. tropicalis*. The overlap of differentially expressed genes between the two *C. albicans* isolates was 48%. This result is consistent with previous work showing that clinical isolates differ in their biofilm forming abilities (Hirakawa *et al*. 2014; Huang *et al*. 2019). As noted in the discussion, we believe that many of the differences among clinical strains arose after the *C. albicans-C. dubliniensis* branch.

To test whether the large differences in the gene expression profiles were specific to the media conditions used, we also performed the transcription profiles in different media, namely Spider for *C. albicans* and *C. dubliniensis*, and RPMI for *C. albicans* and *C. tropicalis*. In the alternative media, the conservation between the sets of genes that changed their expression at least twofold during biofilm formation are even lower, 22% and 11%, respectively for *C. albicans* and *C. dubliniensis*, and *C. albicans* and *C. tropicalis*. Despite these major differences, all the interspecific pairwise overlaps are greater than would be expected by chance (hypergeometric test, *P* << 0.5). Overall, the low degree of conservation in genome-wide gene expression, agrees well with the low conservation of regulator-target gene connections across species described above.

### Analysis of an interspecies hybrid independently supports the conclusions from the species-to-species comparisons

Many challenges exist in mapping and comparing regulator-target gene connections in transcription networks between yeast species and, more generally, between any species (Chen *et al*. 2012). These difficulties include technical issues such as differential nucleic acid recovery and signal to noise ratios, which can vary considerably from one species to the next. However, probably the most difficult problem to circumvent arises from different species having different physiological responses to the same external environment. For example, 30°C could be the optimal temperature for one species but might induce a stress response in a closely related species. Therefore, a network comparison between these species at 30°C might be dominated, not by evolutionary changes in the transcription circuitry *per se*, but simply by the fact that only one species has induced a stress response. This problem can be overcome by creating and analysing interspecies hybrids, where the genomes of two different species are present in the same cell and thus exposed to the same physiological state (Wilson *et al*. 2008; Nocedal *et al*. 2017). This approach, which can only be carried out between closely related species, specifically reveals the *cis*-regulatory changes that have accumulated between the two genomes since the species last shared a common ancestor.

We took advantage of the fact that it is possible to mate *C. albicans* and *C. dubliniensis* (each diploid) to generate tetraploid hybrids (Pujol *et al*. 2004). These hybrids form biofilms similar to those formed by *C. albicans* (Sup Fig 6). We performed ChIP-seq of Ndt80 in this hybrid, immunoprecipitating the *C. albicans* Ndt80 protein in one set of experiments and the *C. dubliniensis* Ndt80 in another set. The results showed that — in the hybrid — the target genes bound by the *C. albicans* Ndt80 and the *C. dubliniensis* Ndt80 were highly correlated, similar to two biological replicates carried out in the same species (Figure 6A). In other words, we obtained the same target genes in the hybrid regardless of which Ndt80 was tagged for immunoprecipitation. Importantly, the binding positions on the *C. dubliniensis* genome in the hybrid were characteristic of the results in *C. dubliniensis*, specifically 97% of the targets in the hybrid are targets in *C. dubliniensis*, and the positions on the *C. albicans* genome were characteristic of *C. albicans* with 96% of the targets in the hybrid being targets in *C. albicans*. Although we only carried out this experiment with one master regulator, the results independently validate our earlier conclusions based on the much more extensive species-to-species comparisons. These observations confirm our previous conclusion that the extreme differences in regulator-target gene connections observed across *Candida* species are due to changes in the *cis*-regulatory sequences in the target genes rather than changes in the regulators themselves or differences in the physiological state of the species at the time of analysis.

**Figure 6.**
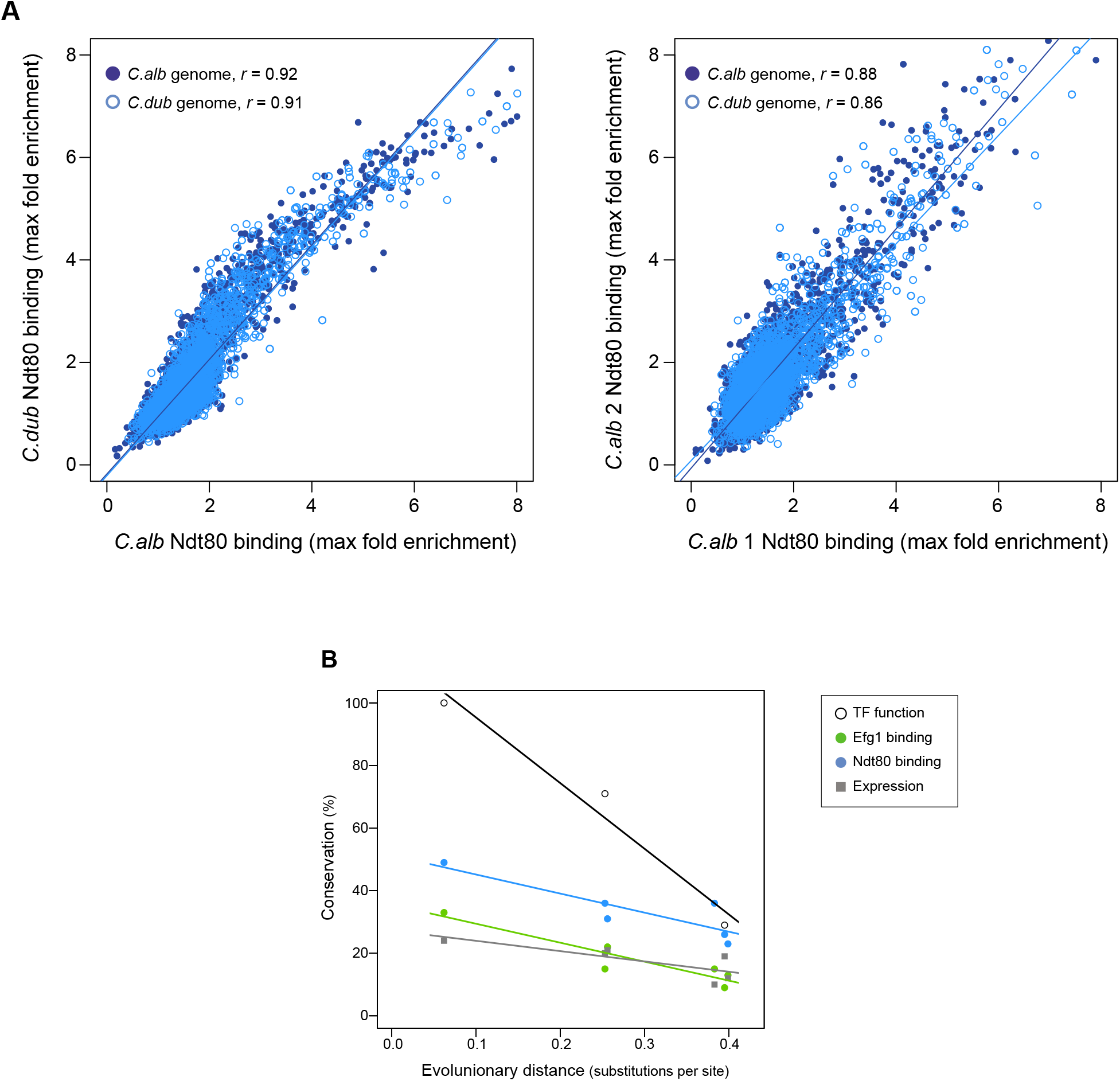
Ndt80 ChIP-seq in a hybrid and rate of conservation change of the different network components. (A) Genome-wide comparison of *C. albicans* and *C. dubliniensis* Ndt80 binding in the hybrid strain. Binding to both the *C. albicans* (dark blue filled dots) and the *C. dubliniensis* (light blue empty dots) genomes is depicted. The maximal fold enrichment for each upstream intergenic region in the genome is plotted as well as the linear regression for each comparison. The left panel shows the *C. albicans* Ndt80–*C. dubliniensis* Ndt80 comparison while the right panel shows, as a reference, the comparison of the two experimental replicates that are most dissimilar. (B) Correlation between the master regulators required for biofilm formation, the Efg1 and Ndt80 binding targets, and biofilm gene expression as a function of evolutionary distance. Master regulator conservation is depicted as the percentage of *C. albicans* regulators required for biofilm formation. Efg1 and Ndt80 target conservation reflect the percentage of targets shared by the different species pairs. Gene expression conservation represents the number of genes whose expression changes at least 1.5 log2 folds under biofilm forming conditions between each species pair. The *C. parapsilosis* gene expression data is from (Holland *et al*. 2014). Linear regressions are shown in the corresponding color. Evolutionary distance as substitutions per site was calculated from a phylogenetic tree of these species, inferred from protein sequences of 73 highly conserved genes (Lohse *et al*. 2013).

## DISCUSSION

In this paper, we examined how a complex transcriptional network underlying a specific phenotype (Figure 1) evolved over a span of approximately 70 million years. The phenotype is biofilm formation by *Candida* species, a group of fungi that colonize humans, sometimes leading to disease. We documented phenotypic differences in biofilm formation across many fungal species, and we mapped the transcriptional networks underlying biofilm formation in four of them, *C. albicans, C. dubliniensis, C. tropicalis* and *C. parapsilosis*. All four species form complex biofilms both *in vitro* and *in vivo* in a rat catheter model (Figure 2).

Using *C. albicans* as a reference species, our analysis leads to the following five main conclusions. (1) as we move away from *C. albicans*, biofilms become less complex, both in terms of structure and of composition; that is, fewer cell types are involved and the resulting biofilm is less regular. At larger evolutionary distances, fungal species did not form biofilms at all under the conditions we tested. (2) Of the seven master transcriptional regulators of biofilm formation in *C. albicans*, all seven are needed in the most closely related species (*C. dubliniensis*), but, as we move further away evolutionarily, fewer are required for biofilm formation. For example, two of the master regulators (Rob1 and Flo8) are dispensable for biofilm formation in the next most closely related species (*C. tropicalis*), and three of the seven are not required in *C. parapsilosis*. As shown by Holland and colleagues (Holland *et al*. 2014), other transcriptional regulators (present in *C. albicans* but not required for biofilm formation in this species) have assumed the role of master regulators in *C. parapsilosis*. (3) In contrast to the relatively slow evolutionary substitutions of master regulators, the connections between the master regulators and their target genes have changed very rapidly over evolutionary time (Figure 6B). This conclusion is most obvious when we compare the two most closely related species, *C. albicans* and *C. dubliniensis*, estimated to have last shared a common ancestor 20 million years ago. Depending on the regulator, fewer than 50% of the master regulator-target gene connections were observed to be conserved. This conclusion was independently verified for one regulator — Ndt80 — by analyzing its binding distribution across the two genomes in a *C. albicans-C. dubliniensis* hybrid; here, the binding distribution of Ndt80 across one genome differed considerably from that of the other, and each resembled that seen in the cognate individual species. This result strongly supports the conclusion that the differences in regulator-target gene connections across species are due largely to changes in the *cis*-regulatory sequences of the target genes rather than changes in the regulators. (4) As predicted from the extensive changes in regulator-target gene connections, mRNA expression during biofilm formation differs considerably from one species to the next. Like the other changes we have documented in this paper, mRNA expression divergence becomes greater as the phylogenetic distance increases (Figure 6). (5) Despite the extensive changes in the transcription networks underlying biofilm formation across the species we examined, several key features of the overall architecture of the network appear to be preserved. For example, all species show high connectivity in the sense that many target genes are directly connected (by binding) to more than one master regulator. Moreover, many of the master regulators bind to their own control regions as well as those of the other master regulators. We have argued elsewhere that these two features are likely to be common to many complex transcription networks (Sorrells and Johnson 2015), and the results presented here show that, despite many changes in individual regulator-target connections, the basic “structural features” of the network are preserved across the biofilm networks of the four species examined.

To place our findings in context it is also instructive to compare our analyses of biofilm formation across species with recent studies where biofilm formation has been analyzed across different isolates of a single species, *C. albicans* (Hirakawa *et al*. 2014; Huang *et al*. 2019). Hirakawa *et al*. (2014) determined the genome sequences of 21 clinical isolates of *C. albicans* and examined their abilities to form biofilms. The genome comparisons revealed many differences among strains including aneuploidies, losses of heterozygosity, and mutations in coding sequences; moreover, the strains differed substantially in their abilities to form biofilms. Among the strains analyzed, the one used in our study (SC5314) was among the thickest biofilm producers as assayed by dry weight; the majority of isolates formed thinner biofilms. One clinical isolate that formed very poor biofilms was found to have an inactivating mutation in *EFG1*, one of the biofilm master regulators, indicating a relatively recent change. Because the *C. dubliniensis* strain used in our study (CD36) formed biofilms that are similar to those of strain SC5314, we believe that SC5314 is a good representative of *C. albicans* and that most of the clinical isolates probably acquired mutations (including aneuploidies and losses of heterozygosity) relatively recently. Huang *et al*. (2019) examined five of the previously sequenced strains in much more detail including the dependence on individual transcriptional regulators for biofilm formation. Although the magnitude of the effect of transcriptional regulator deletions on biofilm formation varied across strains, SC5314 again appears to be a good representation of the ability of *C. albicans* as a species to form thick, complex biofilms.

To our knowledge, this study is the first to examine in detail how a complex transcription network changes over a relatively short evolutionary time — 70 million years — represented by four different species. During that time, the master transcription regulators controlling biofilm formation have slowly changed, but their connections to the target genes they control have changed rapidly. We do not know which, if any, of these changes were adaptive; in this regard, it is important to note that, although the biofilms produced by the *Candida* species have many similarities, they do differ from species to species in at least subtle aspects. Even considering the two most closely related species (*C. albicans* and *C. dubliniensis*), it is possible to distinguish their biofilms. Although they appear very similar under the confocal microscope, the *C. albicans* biofilms form faster and under a greater range of conditions; once formed, they are more difficult to disrupt than those of *C. dubliniensis*. Although these differences may help to explain why *C. albicans* is a greater problem in the clinic than *C. dubliniensis*, it is difficult to reconcile these subtle differences in phenotype with the large differences in the underlying transcriptional circuitry. Given the large magnitude of changes underlying such similar phenotypic output, we propose as a default hypothesis that many of the changes in transcription circuitry result from neutral evolution, more specifically, constructive neutral evolution whereby molecular complexity can change without an increase in fitness (Stoltzfus 1999; Lynch 2007; Wagner 2014; Sorrells and Johnson 2015; Brunet and Doolittle 2018). This study clearly shows that complex transcription networks responsible for the same basic phenotype can undergo evolutionary changes that appear much greater in magnitude than the resulting differences in phenotype.

## METHODS

### Characterization of biofilm formation

Visualization of biofilms by confocal scanning laser microscopy (CSLM) of the different species and strains was performed on silicon squares as described previously (Nobile *et al*. 2012). The strains used for each species and media employed are shown in Sup Table 4 and 1, respectively. Briefly, for the adhesion phase, silicone squares pre-treated with adult bovine serum albumin (BSA) were inoculated to an OD_600_ of 0.5 with cells from an overnight culture grown at 30°C in YPD medium. After incubation for 90 minutes at 37°C and 200 rpm in the specific medium (Sup Table 1) for adhesion, the squares were washed with phosphate-buffered saline (PBS) and then placed in fresh media and incubated for 48 hours at 37°C and 200 rpm. Biofilms of the species that do not grow well at 37°C were grown at 30°C as indicated in Figure 2. After 48 hours, the biofilms were stained for 1 hr with 50 mg/mL of concanavalin A-Alexa Fluor 594 conjugate and visualized on a Nikon Eclipse C1si upright spectral imaging confocal microscope using a 40x/0.80W Nikon objective.

Visualization of biofilm formation over time was also performed by CSLM in Spider medium. For each of the four species observed (*C. albicans, C. dubliniensis, C. tropicalis* and *C. parapsilosis*), seven independent silicone squares were used for biofilm formation as described above and the biofilms at each square were visualized at 30 minutes, 4, 8, 12, 24, 48 and 96 hours after the adhesion phase (Sup Figure 1).

Determination of the biomass dry-weight of biofilms of the different species and strains was performed by growing biofilms on the bottoms of 6-well polystyrene plates pre-treated with BSA as previously described (Nobile *et al*. 2012). The silicone squares were adhered for 90 minutes. These assays were performed in a modified Spider medium that contained 1% glucose rather than mannitol as the carbon source. After 48 hours of biofilm formation, supernatants were aspirated and biofilms were scraped and placed to dry on top of a filter paper. Dried biofilms were weighed on an analytical scale subtracting the weight of a filter paper in which the media without cells was filtered. Five technical replicates were performed per strain. As was performed for CSLM visualization, strains that do not grow well at 37°C were grown at 30°C.

Biofilm formation in a Bioflux microfluidic device (Fluxion Biosciences) was assayed as described previously (Gulati *et al*. 2017). The medium used was Spider with 1% glucose and without mannitol, and assays were performed at 37°C and 30°C for 24 hours.

*In vivo* biofilm formation assays were performed using the rat central-venous catheter infection model as previously described (Andes *et al*. 2004). After 24 hours of infection by the four species tested, biofilm formation on the intraluminal surface of the catheters was observed by scanning electron microscopy (SEM). Procedures were approved by the Institutional Animal Care and Use Committee (IACUC) at the University of Wisconsin, Madison (protocol MV1947).

### Generation of gene deletion knockout strains

Gene deletion knockout strains were constructed using a similar fusion PCR strategy as that described by Noble and Johnson (2005) employing histidine and leucine auxotrophic strains. Construction of these strains was performed using the *SAT1* flipper strategy as previously described (Mancera *et al*. 2019). All the strains employed and generated in this study are shown in Sup Table 4. In brief, the two alleles of each regulator in *C. dubliniensis* were subsequently deleted using the *C. albicans HIS1* and *LEU2* genes. In *C. tropicalis* the first allele was deleted using the *C. albicans LEU2* gene while the second was deleted using the *CaHygB* gene that confers resistance to hygromycin B (Basso *et al*. 2010). To generate the gene-deletion cassettes, ~350 bp flanking 5’ and 3’ regions of each regulator where PCR amplified from genomic DNA, and fused to the corresponding auxotrophic/drug resistance marker by fusion PCR. Transformation was performed by electroporation as previously described (Porman *et al*. 2011). Verification of correct integration of the gene deletion cassettes was performed by colony PCR with primers directed to both flanks of the disrupted gene. Final gene-deletion confirmation was performed by colony PCR with primers that anneal at the ORF of each regulator. Two independent isolates of each deletion mutant originating from two separate transformations were generated for each regulator deletion. The regulator knockout strains of *C. albicans* and *C. parapsilosis* had been previously generated as part of efforts to generate collections of regulator gene knockout strains (Homann *et al*. 2009; Holland *et al*. 2014).

### Chromatin Immunoprecipitation followed by sequencing (ChIP-seq)

Chromatin immunoprecipitation followed by sequencing (ChIP-seq) to identify the target genes of the seven regulators was performed as previously described (Hernday *et al*. 2010; Lohse and Johnson 2016), and sequenced using Illumina HiSeq 2500 or 4000 platforms. Each of the seven regulators in *C. dubliniensis* and *C. tropicalis* was tagged in the wildtype strain background with a 13x Myc epitope tag at the C-terminus from the pADH34 or pEM019 plasmids, respectively, as previously described (Hernday *et al*. 2010; Mancera *et al*. 2019). *C. albicans* Myc-tagged strains had been similarly generated previously (Nobile *et al*. 2012). *C. parapsilosis* Brg1, Ndt80 and Tec1 were tagged using a 6x C-terminal Myc tag amplified from plasmid pFA-MYC-HIS1 as described by (Connolly *et al*. 2013). *C. parapsilosis* Efg1 had been previously tagged (Connolly *et al*. 2013). We were not successful tagging Bcr1 in this species. Genotype details for all the strains generated and used are in Sup Table 4.

To generate the *C. albicans/C. dubliniensis* tagged hybrid strains, the α or **a** allele of the mating-type-like (MTL) locus was deleted in the Ndt80 tagged strains described above. These deletions allowed the strains to become capable of white-opaque switching, and thus mating competent. In *C. albicans*, the α allele of the *MTL* locus was deleted by replacing it with *ARG4* using the plasmid pJD1 as previously described (Lohse *et al*. 2016). In *C. dubliniensis*, the **a** allele of the *MTL* locus was deleted using a cassette containing the *SAT1* nourseothricin resistance marker from plasmid pSFS2A flanked by ~300 bp homology regions identical to the 3’ and 5’ upstream/downstream regions of the *MTL* locus. pSFS2A is a plasmid derived from pSFS2 (Reuss *et al*. 2004) that contains the *SAT1* reusable cassette in the backbone of vector pBC SK+ instead of pBluescript II KS and that was kindly provided by Joachim Morschhauser (U. Würzburg). The deletion of the α or **a** alleles was verified by colony PCR of the two flanks. Generation of the hybrid strains was done by overlaying the mating competent wildtype and Myc-tagged strains on a YPD plate for 48 hours at 30°C. Single colonies were then streaked out and hybrids were selected by growing on media containing nourseothricin and lacking arginine. The hybrid strains were further verified measuring DNA content using FACS. As controls, the two wildtype untagged mating competent strains were hybridized.

All immunoprecipitation experiments were performed under biofilm growth conditions in 6-well polystyrene plates as previously described (Nobile *et al*. 2012). After 48 hours of biofilm growth in Spider 1% glucose at 37°C and 200 rpm, cells were fixed with 1% formaldehyde for 15 minutes. Cell disruption and immunoprecipitation were performed as previously described (Hernday *et al*. 2010) using a c-Myc tag monoclonal antibody (RRID: AB_2536303). After crosslink reversal, instead of performing a phenol/chloroform extraction, we used a MiniElute QIAGEN kit to purify the immunoprecipitated DNA. Library preparation for Illumina sequencing was performed using an NEBNext ChIP-Seq Library Prep Master Mix Set for Illumina sequencing. Between 12 and 24 samples were multiplexed per lane. As controls, immunoprecipitations were performed in matched strains that lacked the Myc tag. Two biological replicates were performed for each regulator in the four species.

### Identification of regulator directly bound target genes by ChIP-seq

ChIP-seq reads were mapped to their corresponding genome using Bowtie 2 with default parameters (Langmead and Salzberg 2012). The genome sequences and annotations were obtained from CGD for versions: 06-Apr-2014C_albicans_SC5314_A21, 06-Nov-2013C_dubliniensis_CD36, 11-Dec-2013C_tropicalis_MYA-3404, and C_parapsilosis_CDC317_version_s01-m03-r13. The SAMtools package was used to convert, sort and index the sequenced reads to BAM format (Li *et al*. 2009). We observed that the peak calling algorithm was more specific and sensitive if the number of reads in the treatment and control datasets were similar. Therefore, we adjusted the number of reads in the different treatment-control dataset pairs using SAMtools view -s function prior to peak calling. Peak calling was performed using MACS2 (Zhang *et al*. 2008) with a q-value cutoff of 0.01; the shiftsize parameter was determined using the SPP package in R (Kharchenko *et al*. 2008). Peaks were considered as true binding events only if the peak was identified in both biological replicates. Assignment of peaks to ORFs was done using MochiView when the peak was present in the intergenic region immediately upstream of the ORF (Homann and Johnson 2010).

To identify regulator binding target genes in the hybrid strains, ChIP-seq reads were aligned to the *C. albicans* and *C. dubliniensis* genomes as described above. Reads that aligned to both genomes were subsequently filtered out. Further processing, peak calling, and assignment of peaks to ORFs were then performed independently for reads that mapped to the *C. albicans* and *C. dubliniensis* genomes as described above.

### *De novo* sequence motif discovery and enrichment for the regulators

DNA-binding motifs were generated *de novo* for regulators from ChIP-Seq experiments using DREME (Bailey 2011). The union of the sequences under the peaks of the two biological replicates for each experiment were tested against a background of matched length random genomic sequences from that species. The top scoring motif was taken and is shown in Figure 5.

Based on the motifs generated *de novo* using DREME as well as previously reported DNA-binding motifs (Lassak *et al*. 2011; Nobile *et al*. 2012; Connolly *et al*. 2013; Nocedal *et al*. 2017), a high-confidence “consensus motif” was generated for Ndt80 (CACAAA) and Efg1 (TGCAT). To determine enrichment of these consensus motifs in peaks identified for each ChIP-Seq experiment, the number of consensus motifs in the union of the sequence under the peaks of both biological replicates was compared to the number of motifs in intergenic regions for that species. A Fisher’s one-tailed exact test was performed to generate a p-value representing enrichment of the motif in peaks compared to matched length random intergenic sequences.

### Genome-wide transcription profiling

Cultures for the extraction of total RNA under biofilm growth conditions were performed on biofilms grown on the bottom of 6-well polystyrene plates for 48 hours at 37°C and 200 rpm as previously described for the determination of biofilm biomass dry-weight (Nobile *et al*. 2012). The media used was Spider 1% glucose for all species, Spider for *C. albicans* and *C. dubliniensis*, and RPMI 1% glucose for *C. albicans* and *C. tropicalis*. Planktonic cultures for total RNA were grown in the corresponding media by inoculating with cells from an overnight 30°C YPD culture to an OD_600_ of 0.05. Cultures were then grown in flasks at 37°C with shaking at 225 rpm until they reached an OD_600_ of 1.0. Biofilm and planktonic cultures were harvested immediately by centrifugation at 3000 g for three minutes and snap-frozen in liquid nitrogen. Total RNA was extracted from the frozen pellets using the RiboPure-Yeast RNA kit (Ambion, AM1926) following the manufacturer’s recommendations. Transcription profiling was performed by hybridization to custom-designed Agilent 8*15k oligonucleotide microarrays that contain between 2 and 3 independent probes for each ORF (*C. albicans* AMADID #020166; *C. dubliniensis*, AMADID #042592; *C. tropicalis* AMADID #042593).

cDNA synthesis, dye coupling, hybridization and microarray analysis was performed as previously described (Nobile *et al*. 2012). Two biological replicates were performed for each species in each condition using the wildtype strains.

## DATA DEPOSITION

ChIP-Seq and microarray gene expression data has been deposited to the NCBI Gene Expression Omnibus (GEO) repository under Superseries GSE160783.

## ACKNOWLEDGMENTS

We would like to thank Derek Sullivan and especially Joachim Morschhäuser for providing strains, plasmids and advice, and Victor Hanson-Smith for help with the ChIP-seq data analysis. This work was supported by a Human Frontier Science Program grant to EM, a UC-MEXUS grant to EM, National Institutes of Health (NIH) grants Ro1AI083311 and Ro1AI049187 to ADJ, Ro1AI073289 to DRA, and R35GM124594 and R21AI125801 to CJN, by a Pew Biomedical Scholar Award from the Pew Charitable Trusts to CJN, the Kamangar family in the form of an endowed chair to CJN, CONACyT grant CB-2016-01 282511 to EM, and a Welcome Trust Seed Award in Science to EM.

## CONTRIBUTIONS

Conceptualization, EM and ADJ; Formal Analysis, EM and IN; Investigation, EM, IN, SH, MG, KFM; Resources, EM, DRA, CJN, GB and ADJ; Writing, EM and ADJ; Supervision, EM, DRA, CJN, GB and ADJ.

---

**Supplementary Table 1**. Biofilm formation of *C. albicans* and its three most closely related species in different media. Biofilms were grown on silicone squares at 37°C with shaking at 200 rpm. Biofilm formation was assessed by CSLM after 48 hours of growth. Spider medium was prepared with 1% nutrient broth, 0.4% potassium phosphate and adjusted to pH 7.2. RPMI media used was RMPI 1640 with 165 mM MOPS and L-glutamine and without sodium bicarbonate (Lonza, 04-525F). YNB media was prepared with 0.67% yeast nitrogen base with ammonium sulfate.

**Supplementary Table 2**. Number of binding regions identified for each regulator in the four *Candida* species studied. “NA”, indicates experiments that were not performed.

**Supplementary Table 3**. Master regulator binding to each other’s upstream intergenic regions. Numbers represent the number of species in which binding was observed. The maximum possible number of species for which we have data is shown in parenthesis for each master regulator.

**Supplementary Table 4**. Strains used in this study.

**Supplementary Figure 1**. Time course of biofilm formation of *C. albicans* and its three most closely related *Candida* species. Biofilms were grown in Spider medium on silicone squares at 37°C with shaking at 200 rpm. Biofilm formation was assessed by CSLM after 30 minutes, 4, 8, 12, 24, 48 and 96 hours of biofilm growth. In each micrograph, the lower panel shows the lateral maximal intensity projection while the upper panel shows the top maximal intensity projection. Scale bars represent 50 μm.

**Supplementary Figure 2**. Biofilm formation in a microfluidic device by different CTG clade species was assayed as previously described (Gulati *et al*. 2017). The medium used was Spider with 1% glucose and without mannitol. Assays were performed at 37°C and 30°C for 24 hours. The hashtag indicates the species that does not grow well at 37°C and for which biofilms were grown at 30°C

**Supplementary Figure 3**. Morphology of biofilms formed by the gene-deletion mutants of the biofilm regulators in *C. dubliniensis* and *C. tropicalis* visualized by CSLM. Biofilms were grown as described in the methods on the surface of silicone squares in Spider medium for *C. dubliniensis* and RPMI medium for *C. tropicalis* for 48 hrs at 37°C. Scale bars represent 50 μm. The *C. tropicalis ndt80* mutant as well as the *flo8* mutants for both species were not analyzed by CSLM.

**Supplementary Figure 4**. Efg1 and Ndt80 target gene conservation between species when using different criteria to identify targets. This was performed similar to what was described previously (Nocedal *et al*. 2017). (A) Regulator gene target conservation when only using significant ChIP-seq signal over control in the upstream intergenic region as a criterion to identify gene targets. (B) Regulator gene target conservation when using ChIP-seq signal as in (A), but also filtering to include only those regions that also had a regulator binding motif in the upstream intergenic region. (C) Regulator gene target conservation when using ChIP-seq signal as in (A), but also two-fold gene expression change during biofilm growth conditions to identify target genes. (D) Regulator gene target conservation when using ChIP-seq signal, regulator binding motif presence and gene expression change during biofilm growth conditions to identify target genes. We did not perform gene expression profiling in *C. parapsilosis* and therefore gene expression change could not be used as a criterion for comparison with *C. parapsilosis*.

**Supplementary Figure 5**. Ndt80 binding throughout the genome in biofilms grown using different media. Biofilms were grown on the bottoms of 6-well polystyrene plates for 48 hrs. as described in the methods. Binding was determined by ChIP-seq for biofilms grown in Spider 1% glucose medium for the three species and compared to binding for biofilms grown in Spider medium for *C. albicans* and *C. dubliniensis*, and RPMI medium for *C. tropicalis*. The maximum fold enrichment of each upstream intergenic region in the genome are plotted as well as the linear regression for each comparison. All correlations are highly significant (Pearson’s correlation p-value << 0.001).

**Supplementary Figure 6**. Morphology of biofilms formed by the *C. albicans-C. dubliniensis* tetraploid hybrid (*C. albicans* x *C. dubliniensis*) visualized by CSLM. For comparison, the biofilms formed by *C. albicans, C. dubliniensis* and a *C. albicans* tetraploid (*C. albicans* x *C. albicans*) are also shown. Biofilms were grown as described in the methods on the surface of silicone squares in Spider medium for 48 hrs at 37°C. Scale bars represent 50 μm.

## REFERENCES

Andes, D., J. Nett, P. Oschel, R. Albrecht, K. Marchillo et al., 2004 Development and characterization of an in vivo central venous catheter Candida albicans biofilm model. Infect Immun 72: 6023–6031.

Araujo, D., M. Henriques and S. Silva, 2017 Portrait of Candida Species Biofilm Regulatory Network Genes. Trends Microbiol 25: 62–75.

Bailey, T. L., 2011 DREME: motif discovery in transcription factor ChIP-seq data. Bioinformatics 27: 1653–1659.

Basso, L. R., Jr., A. Bartiss, Y. Mao, C. E. Gast, P. S. Coelho et al., 2010 Transformation of *Candida albicans* with a synthetic hygromycin B resistance gene. Yeast 27: 1039–1048.

Blankenship, J. R., and A. P. Mitchell, 2006 How to build a biofilm: a fungal perspective. Curr Opin Microbiol 9: 588–594.

Brunet, T. D. P., and W. F. Doolittle, 2018 The generality of Constructive Neutral Evolution. Biology & Philosophy 33: 2.

Butler, G., M. D. Rasmussen, M. F. Lin, M. A. Santos, S. Sakthikumar et al., 2009 Evolution of pathogenicity and sexual reproduction in eight Candida genomes. Nature 459: 657–662.

Byrne, K. P., and K. H. Wolfe, 2005 The Yeast Gene Order Browser: combining curated homology and syntenic context reveals gene fate in polyploid species. Genome Res 15: 1456–1461.

Calderone, R. A., and C. J. Clancy, 2012 Candida and Candidiasis, Second Edition. American Society of Microbiology.

Chen, Y., N. Negre, Q. Li, J. O. Mieczkowska, M. Slattery et al., 2012 Systematic evaluation of factors influencing ChIP-seq fidelity. Nat Methods 9: 609–614.

Connolly, L. A., A. Riccombeni, Z. Grozer, L. M. Holland, D. B. Lynch et al., 2013 The APSES transcription factor Efg1 is a global regulator that controls morphogenesis and biofilm formation in Candida parapsilosis. Mol Microbiol 90: 36–53.

Ding, C., and G. Butler, 2007 Development of a gene knockout system in Candida parapsilosis reveals a conserved role for BCR1 in biofilm formation. Eukaryot Cell 6: 1310–1319.

Dominguez, E., R. Zarnowski, H. Sanchez, A. S. Covelli, W. M. Westler et al., 2018 Conservation and Divergence in the Candida Species Biofilm Matrix Mannan-Glucan Complex Structure, Function, and Genetic Control. mBio 9.

Donlan, R. M., 2001 Biofilm formation: a clinically relevant microbiological process. Clin Infect Dis 33: 1387–1392.

Fox, E. P., C. K. Bui, J. E. Nett, N. Hartooni, M. C. Mui et al., 2015 An expanded regulatory network temporally controls Candida albicans biofilm formation. Mol Microbiol 96: 1226–1239.

Gabaldon, T., M. A. Naranjo-Ortiz and M. Marcet-Houben, 2016 Evolutionary genomics of yeast pathogens in the Saccharomycotina. FEMS Yeast Res 16.

Garcia-Sanchez, S., S. Aubert, I. Iraqui, G. Janbon, J. M. Ghigo et al., 2004 Candida albicans biofilms: a developmental state associated with specific and stable gene expression patterns. Eukaryot Cell 3: 536–545.

Gulati, M., C. L. Ennis, D. L. Rodriguez and C. J. Nobile, 2017 Visualization of Biofilm Formation in Candida albicans Using an Automated Microfluidic Device. J Vis Exp.

Hernday, A. D., S. M. Noble, Q. M. Mitrovich and A. D. Johnson, 2010 Genetics and molecular biology in Candida albicans. Methods Enzymol 470: 737–758.

Hirakawa, M. P., D. A. Martinez, S. Sakthikumar, M. Anderson, A. Berlin et al., 2014 Genetic and phenotypic intra-species variation in *Candida albicans*. Genome Research.

Holland, L. M., M. S. Schroder, S. A. Turner, H. Taff, D. Andes et al., 2014 Comparative phenotypic analysis of the major fungal pathogens Candida parapsilosis and Candida albicans. PLoS Pathog 10: e1004365.

Homann, O. R., J. Dea, S. M. Noble and A. D. Johnson, 2009 A phenotypic profile of the Candida albicans regulatory network. PLoS Genet 5: e1000783.

Homann, O. R., and A. D. Johnson, 2010 MochiView: versatile software for genome browsing and DNA motif analysis. BMC Biol 8: 49.

Huang, M. Y., C. A. Woolford, G. May, C. J. McManus and A. P. Mitchell, 2019 Circuit diversification in a biofilm regulatory network. PLoS Pathog 15: e1007787.

Johnson, D. S., A. Mortazavi, R. M. Myers and B. Wold, 2007 Genome-wide mapping of in vivo protein-DNA interactions. Science 316: 1497–1502.

Kharchenko, P. V., M. Y. Tolstorukov and P. J. Park, 2008 Design and analysis of ChIP-seq experiments for DNA-binding proteins. Nat Biotechnol 26: 1351–1359.

Krassowski, T., A. Y. Coughlan, X. X. Shen, X. Zhou, J. Kominek et al., 2018 Evolutionary instability of CUG-Leu in the genetic code of budding yeasts. Nat Commun 9: 1887.

Kucharikova, S., H. Tournu, K. Lagrou, P. Van Dijck and H. Bujdakova, 2011 Detailed comparison of Candida albicans and Candida glabrata biofilms under different conditions and their susceptibility to caspofungin and anidulafungin. J Med Microbiol 60: 1261–1269.

Kullberg, B. J., and M. C. Arendrup, 2015 Invasive Candidiasis. N Engl J Med 373: 1445–1456.

Kumari, A., S. Mankotia, B. Chaubey, M. Luthra and R. Singh, 2018 Role of biofilm morphology, matrix content and surface hydrophobicity in the biofilm-forming capacity of various Candida species. J Med Microbiol 67: 889–892.

Kurtzman, C. P., J. W. Fell and T. Boekhout, 2011 The Yeasts.

Langmead, B., and S. L. Salzberg, 2012 Fast gapped-read alignment with Bowtie 2. Nat Methods 9: 357–359.

Lassak, T., E. Schneider, M. Bussmann, D. Kurtz, J. R. Manak et al., 2011 Target specificity of the Candida albicans Efg1 regulator. Molecular Microbiology 82: 602–618.

Li, H., B. Handsaker, A. Wysoker, T. Fennell, J. Ruan et al., 2009 The Sequence Alignment/Map format and SAMtools. Bioinformatics 25: 2078–2079.

Lohse, M. B., I. V. Ene, V. B. Craik, A. D. Hernday, E. Mancera et al., 2016 Systematic Genetic Screen for Transcriptional Regulators of the Candida albicans White-Opaque Switch. Genetics 203: 1679–1692.

Lohse, M. B., M. Gulati, A. D. Johnson and C. J. Nobile, 2018 Development and regulation of single- and multi-species Candida albicans biofilms. Nat Rev Microbiol 16: 19–31.

Lohse, M. B., M. Gulati, A. Valle Arevalo, A. Fishburn, A. D. Johnson et al., 2017 Assessment and Optimizations of Candida albicans In Vitro Biofilm Assays. Antimicrob Agents Chemother 61.

Lohse, M. B., A. D. Hernday, P. M. Fordyce, L. Noiman, T. R. Sorrells et al., 2013 Identification and characterization of a previously undescribed family of sequence-specific DNA-binding domains. Proc Natl Acad Sci U S A 110: 7660–7665.

Lohse, M. B., and A. D. Johnson, 2016 Identification and Characterization of Wor4, a New Transcriptional Regulator of White-Opaque Switching. G3 (Bethesda) 6: 721–729.

Lynch, D. P., 1994 Oral candidiasis. History, classification, and clinical presentation. Oral Surg Oral Med Oral Pathol 78: 189–193.

Lynch, M., 2007 The origins of genome architecture. Sinauer Associates Sunderland, MA.

Maguire, S. L., S. S. Oheigeartaigh, K. P. Byrne, M. S. Schroder, P. O’Gaora et al., 2013 Comparative Genome Analysis and Gene Finding in Candida Species Using CGOB. Mol Biol Evol 30: 1281–1291.

Mancera, E., C. Frazer, A. M. Porman, S. Ruiz-Castro, A. D. Johnson et al., 2019 Genetic Modification of Closely Related Candida Species. Front Microbiol 10: 357.

Mishra, P. K., M. Baum and J. Carbon, 2007 Centromere size and position in Candida albicans are evolutionarily conserved independent of DNA sequence heterogeneity. Mol Genet Genomics 278: 455–465.

Moran, G. P., D. C. Coleman and D. J. Sullivan, 2012 Candida albicans versus Candida dubliniensis: Why Is C. albicans More Pathogenic? Int J Microbiol 2012: 205921.

Nobile, C. J., E. P. Fox, J. E. Nett, T. R. Sorrells, Q. M. Mitrovich et al., 2012 A recently evolved transcriptional network controls biofilm development in Candida albicans. Cell 148: 126–138.

Nobile, C. J., and A. D. Johnson, 2015 Candida albicans Biofilms and Human Disease. Annu Rev Microbiol 69: 71–92.

Noble, S. M., and A. D. Johnson, 2005 Strains and strategies for large-scale gene deletion studies of the diploid human fungal pathogen *Candida albicans*. Eukaryot Cell 4: 298–309.

Nocedal, I., E. Mancera and A. D. Johnson, 2017 Gene regulatory network plasticity predates a switch in function of a conserved transcription regulator. Elife 6.

Porman, A. M., K. Alby, M. P. Hirakawa and R. J. Bennett, 2011 Discovery of a phenotypic switch regulating sexual mating in the opportunistic fungal pathogen Candida tropicalis. Proc Natl Acad Sci U S A 108: 21158–21163.

Pujol, C., K. J. Daniels, S. R. Lockhart, T. Srikantha, J. B. Radke et al., 2004 The closely related species Candida albicans and Candida dubliniensis can mate. Eukaryot Cell 3: 1015–1027.

Pujol, C., K. J. Daniels and D. R. Soll, 2015 Comparison of Switching and Biofilm Formation between MTL-Homozygous Strains of Candida albicans and Candida dubliniensis. Eukaryot Cell 14: 1186–1202.

Ramage, G., K. Vande Walle, B. L. Wickes and J. L. Lopez-Ribot, 2001 Biofilm formation by Candida dubliniensis. J Clin Microbiol 39: 3234–3240.

Reuss, O., A. Vik, R. Kolter and J. Morschhauser, 2004 The *SAT1* flipper, an optimized tool for gene disruption in *Candida albicans*. Gene 341: 119–127.

Richard, M. L., C. J. Nobile, V. M. Bruno and A. P. Mitchell, 2005 Candida albicans biofilm-defective mutants. Eukaryot Cell 4: 1493–1502.

Romo, J. A., and C. A. Kumamoto, 2020 On Commensalism of Candida. J Fungi (Basel) 6.

Silva, S., M. Negri, M. Henriques, R. Oliveira, D. W. Williams et al., 2011 Adherence and biofilm formation of non-Candida albicans Candida species. Trends Microbiol 19: 241–247.

Sorrells, T. R., and A. D. Johnson, 2015 Making sense of transcription networks. Cell 161: 714–723.

Stoltzfus, A., 1999 On the possibility of constructive neutral evolution. J Mol Evol 49: 169–181.

Turner, S. A., and G. Butler, 2014 The Candida pathogenic species complex. Cold Spring Harb Perspect Med 4: a019778.

Wagner, A., 2014 Arrival of the fittest: How nature innovates. Current.

Wilson, M. D., N. L. Barbosa-Morais, D. Schmidt, C. M. Conboy, L. Vanes et al., 2008 Species-Specific Transcription in Mice Carrying Human Chromosome 21. Science 322: 434–438.

Zhang, Y., T. Liu, C. A. Meyer, J. Eeckhoute, D. S. Johnson et al., 2008 Model-based analysis of ChIP-Seq (MACS). Genome Biol 9: R137.

