## Supplemental Figures for "Evolution of the complex transcription network controlling biofilm formation in *Candida* species"

Supplementary Figure 1

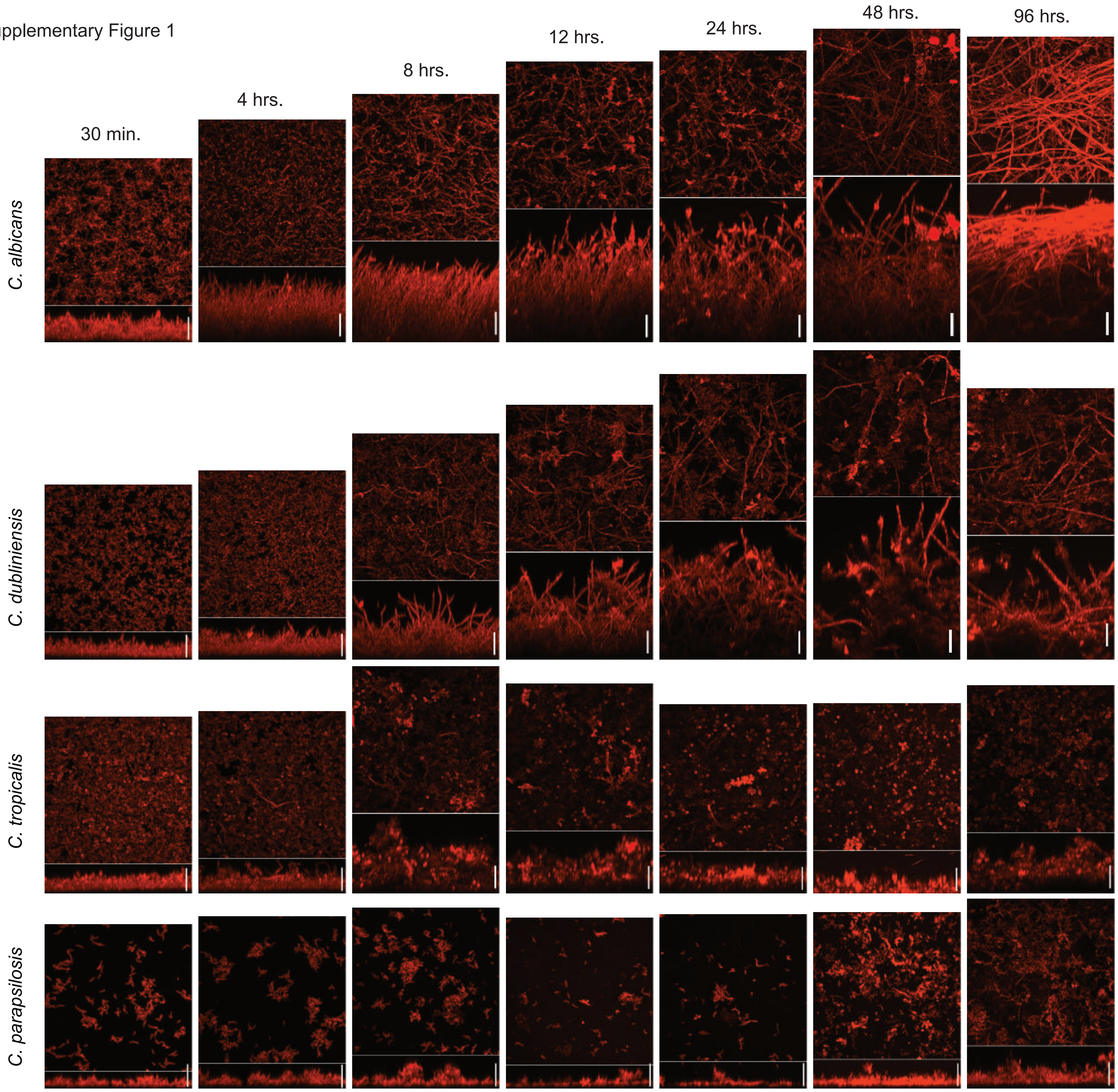

Supplementary Figure 2

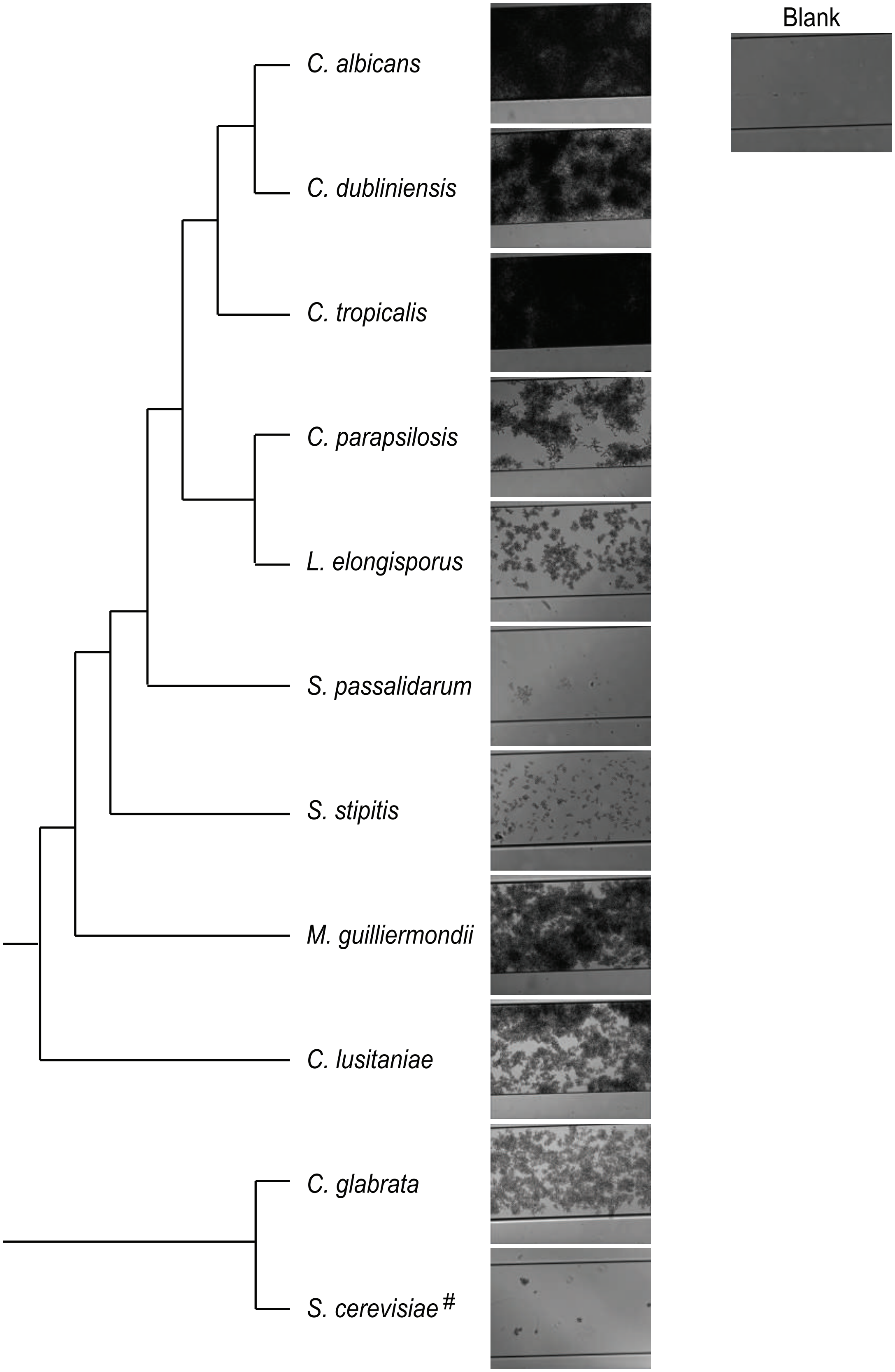

### Supplementary Figure 3

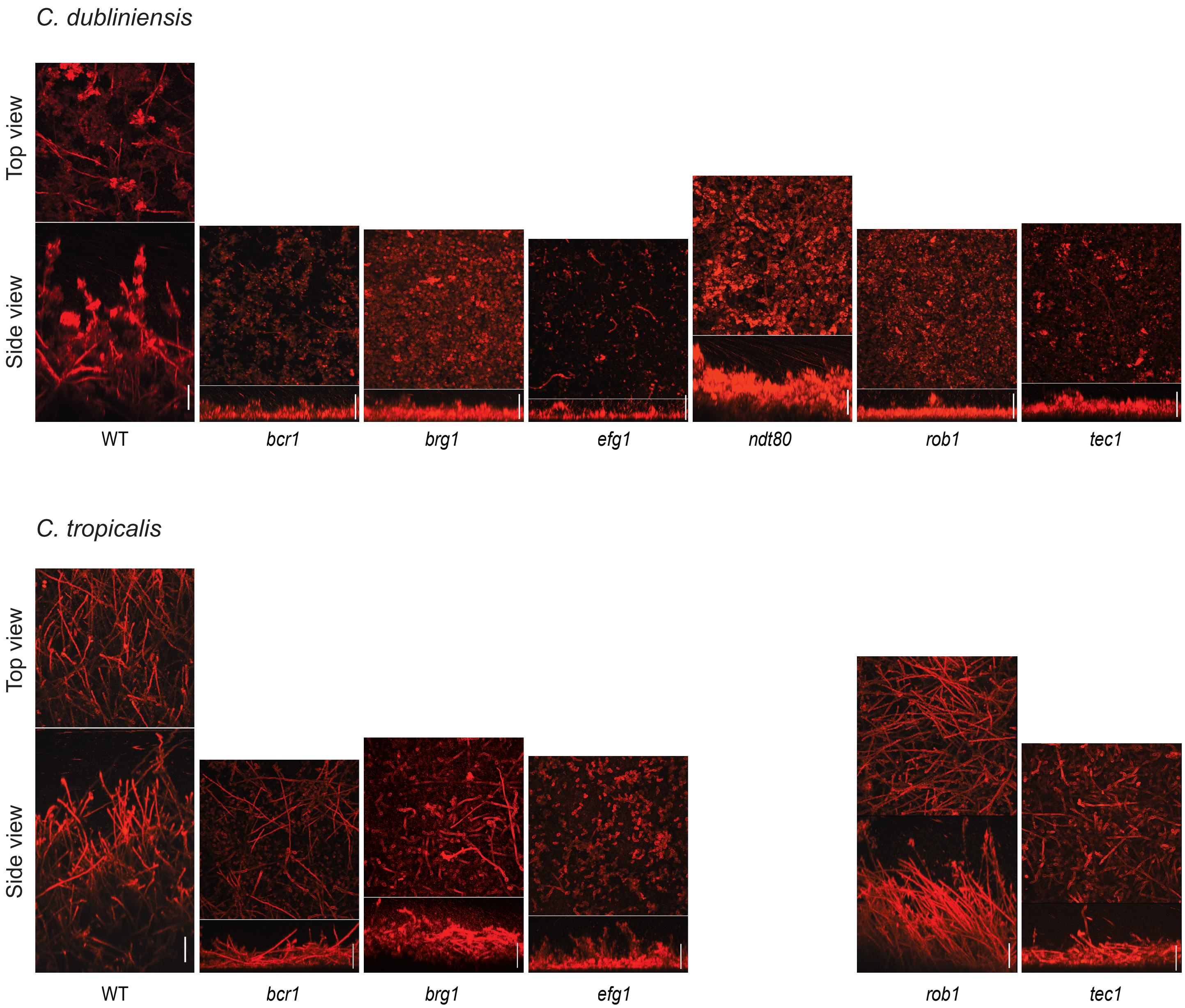

Supplementary Figure 4

**A** **Peak**

**Efg1**

| | *C.alb* | *C.dub* | *C.tro* |
|---|---|---|---|
| *C.dub* | 33 | | |
| *C.tro* | 15 | 22 | |
| *C.par* | 9 | 13 | 15 |

**Ndt80**

| | *C.alb* | *C.dub* | *C.tro* |
|---|---|---|---|
| *C.dub* | 49 | | |
| *C.tro* | 36 | 31 | |
| *C.par* | 26 | 23 | 36 |

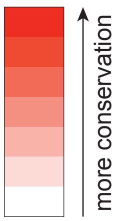


**B** **Peak + motif**

**Efg1**

| | *C.alb* | *C.dub* | *C.tro* |
|---|---|---|---|
| *C.dub* | 32 | | |
| *C.tro* | 12 | 19 | |
| *C.par* | 8 | 12 | 14 |

**Ndt80**

| | *C.alb* | *C.dub* | *C.tro* |
|---|---|---|---|
| *C.dub* | 51 | | |
| *C.tro* | 35 | 31 | |
| *C.par* | 27 | 23 | 34 |

**C** **Peak + expression**

**Efg1**

| | *C.alb* | *C.dub* | *C.tro* |
|---|---|---|---|
| *C.dub* | 19 | | |
| *C.tro* | 12 | 17 | |
| *C.par* | NA | NA | NA |

**Ndt80**

| | *C.alb* | *C.dub* | *C.tro* |
|---|---|---|---|
| *C.dub* | 27 | | |
| *C.tro* | 19 | 16 | |
| *C.par* | NA | NA | NA |

**D** **Peak + motif + expression**

**Efg1**

| | *C.alb* | *C.dub* | *C.tro* |
|---|---|---|---|
| *C.dub* | 19 | | |
| *C.tro* | 11 | 11 | |
| *C.par* | NA | NA | NA |

**Ndt80**

| | *C.alb* | *C.dub* | *C.tro* |
|---|---|---|---|
| *C.dub* | 28 | | |
| *C.tro* | 18 | 15 | |
| *C.par* | NA | NA | NA |

Supplementary Figure 5

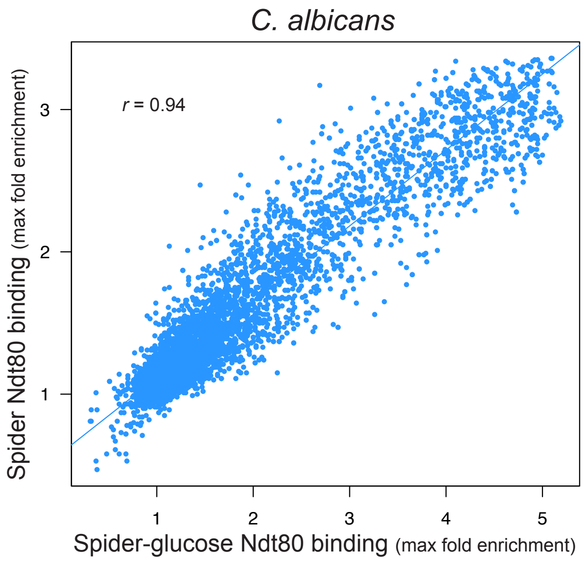


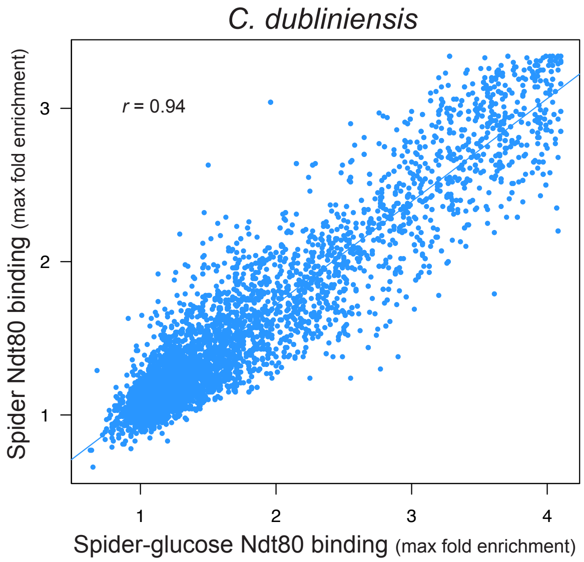


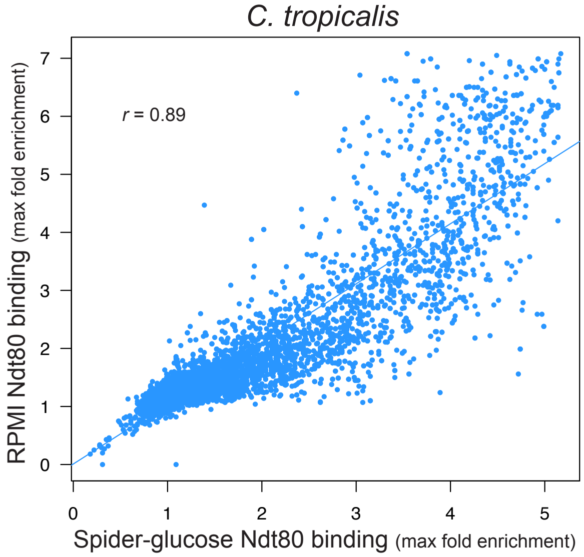

Supplementary Figure 6

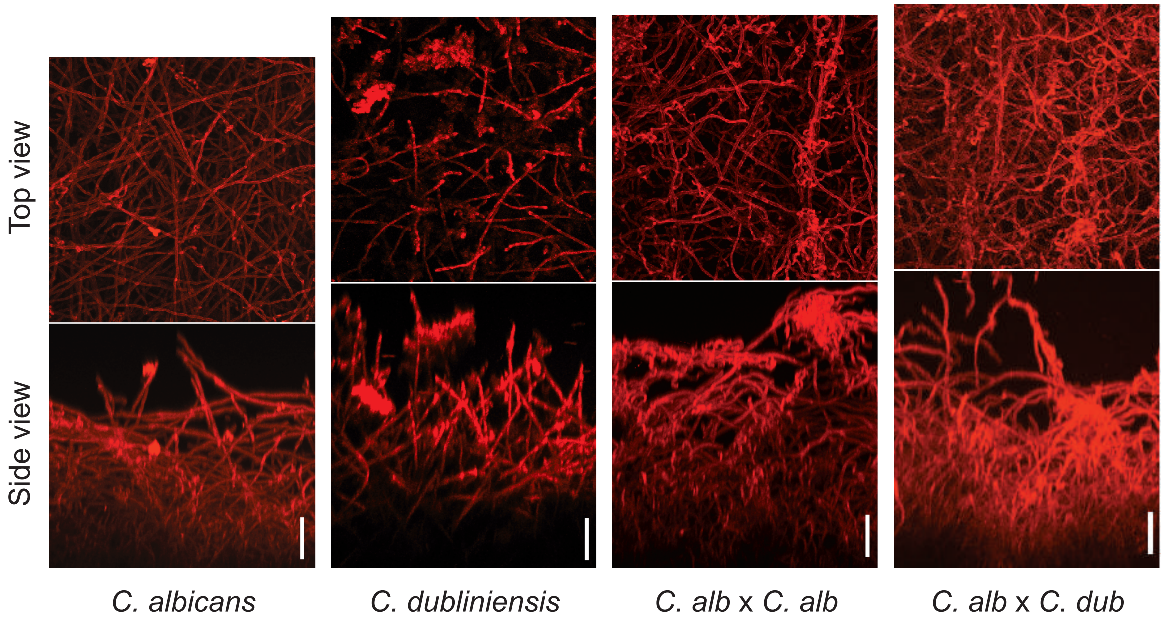
