## Supplemental Tables for "Evolution of the complex transcription network controlling biofilm formation in *Candida* species"

Supplementary Table 1. Biofilm formation of *C. albicans* and its 3 most closely related species in different media.
Biofilms were grown on silicone squares at 37°C with 200 rmp shaking and its formation was assessed by CSLM after 48 hours of growth.

| Media | Carbon source | *C.albicans* | *C.dubliniensis* | *C.tropicalis* | *C.parapsilosis* |
|---|---|---|---|---|---|
| Spider | 1% manitol | + | + | - | - |
| Spider | 1% glucose | ++ | + | + | + |
| Spider | 1% galactose | ++ | + | - | -+ |
| RPMI | 0.2% glucose | + | - | + | - |
| RPMI | 2% glucose | ++ | -+ | + | - |
| YNB | 1% glucose | + | - | ? | + |
| YNB | 2% glucose | + | - | ? | ? |
| YNB | 1% galactose | + | - | ? | ? |

Supplementary Table 2. Number of genes bound by each regulator in the four *Candida* species studied. NA, shows experiments that were not performed.

| Regulator | *C.albicans* | *C.dubliniensis* | *C.tropicalis* | *C.parapsilosis* |
|---|---|---|---|---|
| Bcr1 | 40 | 4 | 10 | NA |
| Brg1 | 98 | 354 | 0 | 0 |
| Efg1 | 168 | 311 | 920 | 609 |
| Flo8 | 0 | 0 | 0 | NA |
| Ndt80 | 1091 | 957 | 1749 | 2927 |
| Rob1 | 19 | 3 | 21 | NA |
| Tec1 | 154 | 0 | 0 | 399 |

Supplementary Table 3. Master regulator binding to each other's upstream intergenic region.
Numbers represent the number of species in which binding was observed.
The maximum possible number of species for which we have data is in parenthesis for each master regulator.

| | | Binding | | | | | |
|---|---|---|---|---|---|---|---|
| | **Name** | **Bcr1 (3)** | **Brg1 (3)** | **Efg1 (4)** | **Ndt80_B (4)** | **Rob1 (3)** | **Tec1 (2)** |
| | **Bcr1** | 1 | 2 | 3 | 4 | 0 | 1 |
| | **Brg1** | 0 | 2 | 3 | 4 | 0 | 2 |
| | **Efg1** | 0 | 1 | 4 | 4 | 1 | 2 |
| **Target** | **Flo8** | 0 | 2 | 1 | 3 | 0 | 0 |
| | **Ndt80_B** | 0 | 0 | 2 | 4 | 0 | 1 |
| | **Rob1** | 0 | 0 | 1 | 2 | 0 | 0 |
| | **Tec1** | 0 | 2 | 3 | 4 | 1 | 0 |

**Supplementary Table 4. Strains used in this study**

| Name | Species | Background | Genotype | Source/Reference |
|---|---|---|---|---|
| SN250 | *C. albicans* | SC5314 | *ura3Δ::λimm434::URA3-IRO1 ura3Δ::λimm434 arg4::hisG arg4::hisG his1::hisG his1::hisG leu2::hisG::CdHIS1 leu2::hisG::CmLEU2* | Noble S.M. *et al*, 2010 |
| CJN1734 | *C. albicans* | SC5314 | *ura3Δ::λimm434::URA3-IRO1 ura3Δ::λimm434 arg4::hisG arg4::hisG his1::hisG his1::hisG leu2::hisG::CdHIS1 leu2::hisG::CmLEU2 BRG1-13XMyc-FRT/BRG1* | Nobile C.J. *et al*, 2012 |
| CJN1748 | *C. albicans* | SC5314 | *ura3Δ::λimm434::URA3-IRO1 ura3Δ::λimm434 arg4::hisG arg4::hisG his1::hisG his1::hisG leu2::hisG::CdHIS1 leu2::hisG::CmLEU2 NDT80-13XMyc-FRT/NDT80* | Nobile C.J. *et al*, 2012 |
| CJN1781 | *C. albicans* | SC5314 | *ura3Δ::λimm434::URA3-IRO1 ura3Δ::λimm434 arg4::hisG arg4::hisG his1::hisG his1::hisG leu2::hisG::CdHIS1 leu2::hisG::CmLEU2 EFG1-13XMyc-FRT/EFG1* | Nobile C.J. *et al*, 2012 |
| CJN1787 | *C. albicans* | SC5314 | *ura3Δ::λimm434::URA3-IRO1 ura3Δ::λimm434 arg4::hisG arg4::hisG his1::hisG his1::hisG leu2::hisG::CdHIS1 leu2::hisG::CmLEU2 BCR1-13XMyc-FRT/BCR1* | Nobile C.J. *et al*, 2012 |
| CJN1865 | *C. albicans* | SC5314 | *ura3Δ::λimm434::URA3-IRO1 ura3Δ::λimm434 arg4::hisG arg4::hisG his1::hisG his1::hisG leu2::hisG::CdHIS1 leu2::hisG::CmLEU2 TEC1-13XMyc-FRT/TEC1* | Nobile C.J. *et al*, 2012 |
| CJN2208 | *C. albicans* | SC5314 | *ura3Δ::λimm434::URA3-IRO1 ura3Δ::λimm434 arg4::hisG arg4::hisG his1::hisG his1::hisG leu2::hisG::CdHIS1 leu2::hisG::CmLEU2 ROB1-7XMyc-FRT/ROB1* | Nobile C.J. *et al*, 2012 |
| CKB36 | *C. albicans* | SC5314 | *ura3Δ::λimm434::URA3-IRO1 ura3Δ::λimm434 arg4::hisG arg4::hisG his1::hisG his1::hisG leu2::hisG::CdHIS1 leu2::hisG::CmLEU2 FLO8-13XMyc-FRT/FLO8* | Nobile C.J. *et al*, 2012 |
| TF022 | *C. albicans* | SC5314 | *arg4Δ/arg4Δ leu2Δ/leu2Δ his1Δ/his1Δ URA3/ura3Δ::imm434 IRO1/iro1Δ::imm434 brg1Δ::HIS1/brg1Δ::LEU2* | Homann O. R., *et al*, 2009 |
| TF095 | *C. albicans* | SC5314 | *arg4Δ/arg4Δ leu2Δ/leu2Δ his1Δ/his1Δ URA3/ura3Δ::imm434 IRO1/iro1Δ::imm434 ndt80Δ::HIS1/ndt80Δ::LEU2* | Homann O. R., *et al*, 2009 |
| TF110 | *C. albicans* | SC5314 | *arg4Δ/arg4Δ leu2Δ/leu2Δ his1Δ/his1Δ URA3/ura3Δ::imm434 IRO1/iro1Δ::imm434 rob1Δ::HIS1/rob1Δ::LEU2* | Homann O. R., *et al*, 2009 |
| TF115 | *C. albicans* | SC5314 | *arg4Δ/arg4Δ leu2Δ/leu2Δ his1Δ/his1Δ URA3/ura3Δ::imm434 IRO1/iro1Δ::imm434 tec1Δ::HIS1/tec1Δ::LEU2* | Homann O. R., *et al*, 2009 |
| TF137 | *C. albicans* | SC5314 | *arg4Δ/arg4Δ leu2Δ/leu2Δ his1Δ/his1Δ URA3/ura3Δ::imm434 IRO1/iro1Δ::imm434 bcr1Δ::HIS1/bcr1Δ::LEU2* | Homann O. R., *et al*, 2009 |
| TF156 | *C. albicans* | SC5314 | *arg4Δ/arg4Δ leu2Δ/leu2Δ his1Δ/his1Δ URA3/ura3Δ::imm434 IRO1/iro1Δ::imm434 efg1Δ::HIS1/efg1Δ::LEU2* | Homann O. R., *et al*, 2009 |
| CD36 | *C. dubliniensis* | CD36 | *WT* | Derek Sullivan |
| CEM091 | *C. dubliniensis* | CD36 | *his1Δ::HIS1 Calb/his1Δ::FRT leu2Δ::LEU2 Calb/leu2Δ::FRT arg4::FRT/arg4::FRT* | Mancera E. *et al*, 2019 |
| CEM100 | *C. dubliniensis* | CD36 | *his1Δ::FRT/his1Δ::FRT leu2Δ::FRT/leu2Δ::FRT arg4::FRT/arg4::FRT bcr1Δ::HIS1 Calb/bcr1Δ::LEU2 Calb* | This study |
| CEM101 | *C. dubliniensis* | CD36 | *his1Δ::FRT/his1Δ::FRT leu2Δ::FRT/leu2Δ::FRT arg4::FRT/arg4::FRT brg1Δ::HIS1 Calb/brg1Δ::LEU2 Calb* | This study |
| CEM102 | *C. dubliniensis* | CD36 | *his1Δ::FRT/his1Δ::FRT leu2Δ::FRT/leu2Δ::FRT arg4::FRT/arg4::FRT efg1Δ::HIS1 Calb/efg1Δ::LEU2 Calb* | This study |
| CEM103 | *C. dubliniensis* | CD36 | *his1Δ::FRT/his1Δ::FRT leu2Δ::FRT/leu2Δ::FRT arg4::FRT/arg4::FRT ndt80Δ::HIS1 Calb/ ndt80Δ::LEU2 Calb* | This study |
| CEM104 | *C. dubliniensis* | CD36 | *his1Δ::FRT/his1Δ::FRT leu2Δ::FRT/leu2Δ::FRT arg4::FRT/arg4::FRT rob1Δ::HIS1 Calb/rob1Δ::LEU2 Calb* | This study |
| CEM105 | *C. dubliniensis* | CD36 | *his1Δ::FRT/his1Δ::FRT leu2Δ::FRT/leu2Δ::FRT arg4::FRT/arg4::FRT tec1Δ::HIS1 Calb/tec1Δ::LEU2 Calb* | This study |
| CEM106 | *C. dubliniensis* | CD36 | *his1Δ::FRT/his1Δ::FRT leu2Δ::FRT/leu2Δ::FRT arg4::FRT/arg4::FRT bcr1Δ::HIS1 Calb/bcr1Δ::LEU2 Calb* | This study |
| CEM107 | *C. dubliniensis* | CD36 | *his1Δ::FRT/his1Δ::FRT leu2Δ::FRT/leu2Δ::FRT arg4::FRT/arg4::FRT brg1Δ::HIS1 Calb/brg1Δ::LEU2 Calb* | This study |
| CEM108 | *C. dubliniensis* | CD36 | *his1Δ::FRT/his1Δ::FRT leu2Δ::FRT/leu2Δ::FRT arg4::FRT/arg4::FRT efg1Δ::HIS1 Calb/efg1Δ::LEU2 Calb* | This study |
| CEM109 | *C. dubliniensis* | CD36 | *his1Δ::FRT/his1Δ::FRT leu2Δ::FRT/leu2Δ::FRT arg4::FRT/arg4::FRT ndt80Δ::HIS1 Calb/ ndt80Δ::LEU2 Calb* | This study |
| CEM110 | *C. dubliniensis* | CD36 | *his1Δ::FRT/his1Δ::FRT leu2Δ::FRT/leu2Δ::FRT arg4::FRT/arg4::FRT rob1Δ::HIS1 Calb/rob1Δ::LEU2 Calb* | This study |
| CEM111 | *C. dubliniensis* | CD36 | *his1Δ::FRT/his1Δ::FRT leu2Δ::FRT/leu2Δ::FRT arg4::FRT/arg4::FRT tec1Δ::HIS1 Calb/tec1Δ::LEU2 Calb* | This study |
| CEM339 | *C. dubliniensis* | CD36 | *his1Δ::HIS1 Calb/his1Δ::FRT leu2Δ::LEU2 Calb/leu2Δ::FRT arg4::FRT/arg4::FRT BCR1/BCR1-13Myc-FRT* | This study |
| CEM340 | *C. dubliniensis* | CD36 | *his1Δ::HIS1 Calb/his1Δ::FRT leu2Δ::LEU2 Calb/leu2Δ::FRT arg4::FRT/arg4::FRT BRG1/BRG1-13Myc-FRT* | This study |
| CEM341 | *C. dubliniensis* | CD36 | *his1Δ::HIS1 Calb/his1Δ::FRT leu2Δ::LEU2 Calb/leu2Δ::FRT arg4::FRT/arg4::FRT EFG1/EFG1-13Myc-FRT* | This study |
| CEM342 | *C. dubliniensis* | CD36 | *his1Δ::HIS1 Calb/his1Δ::FRT leu2Δ::LEU2 Calb/leu2Δ::FRT arg4::FRT/arg4::FRT NDT80/NDT80-13Myc-FRT* | This study |
| CEM343 | *C. dubliniensis* | CD36 | *his1Δ::HIS1 Calb/his1Δ::FRT leu2Δ::LEU2 Calb/leu2Δ::FRT arg4::FRT/arg4::FRT ROB1/ROB1-13Myc-FRT* | This study |
| CEM344 | *C. dubliniensis* | CD36 | *his1Δ::HIS1 Calb/his1Δ::FRT leu2Δ::LEU2 Calb/leu2Δ::FRT arg4::FRT/arg4::FRT TEC1/TEC1-13Myc-FRT* | This study |
| CEM365 | *C. dubliniensis* | CD36 | *his1Δ::FRT/his1Δ::FRT leu2Δ::FRT/leu2Δ::FRT arg4::FRT/arg4::FRT flo8Δ::HIS1 Calb/flo8Δ::LEU2 Calb* | This study |
| CEM366 | *C. dubliniensis* | CD36 | *his1Δ::FRT/his1Δ::FRT leu2Δ::FRT/leu2Δ::FRT arg4::FRT/arg4::FRT flo8Δ::HIS1 Calb/flo8Δ::LEU2 Calb* | This study |
| CEM432 | *C. dubliniensis* | CD36 | *his1Δ::HIS1 Calb/his1Δ::FRT leu2Δ::LEU2 Calb/leu2Δ::FRT arg4::FRT/arg4::FRT FLO8/FLO8-13Myc-FRT* | This study |
| CLIB214 | *C. parapsilosis* | CLIB214 | *WT* | Geraldine Butler |
| CEM385 | *C. parapsilosis* | CLIB214 | *leu2::FRT/leu2::FRT his1::FRT/his1::FRT brg1::LEU2/BRG1::MYC::HIS1* | This study |
| CEM391 | *C. parapsilosis* | CLIB214 | *leu2::FRT/leu2::FRT his1::FRT/his1::FRT efg1::LEU2/EFG1::MYC::HIS1* | This study |
| CEM401 | *C. parapsilosis* | CLIB214 | *leu2::FRT/leu2::FRT his1::FRT/his1::FRT ndt80::LEU2/NDT80::MYC::HIS1* | This study |
| CEM407 | *C. parapsilosis* | CLIB214 | *leu2::FRT/leu2::FRT his1::FRT/his1::FRT tec1::LEU2/TEC1::MYC::HIS1* | This study |
| CEM010 | *C. tropicalis* | MYA3404 | *WT* | ATCC |
| CEM300 | *C. tropicalis* | MYA3404 | *leu2Δ::Calb LEU2/leu2Δ::FRT* | Mancera E. *et al*, 2019 |
| CEM225 | *C. tropicalis* | MYA3404 | *leu2Δ::FRT/leu2Δ::FRT bcr1Δ::LEU2 Calb/bcr1Δ::HYG* | This study |
| CEM226 | *C. tropicalis* | MYA3404 | *leu2Δ::FRT/leu2Δ::FRT bcr1Δ::LEU2 Calb/bcr1Δ::HYG* | This study |
| CEM235 | *C. tropicalis* | MYA3404 | *leu2Δ::FRT/leu2Δ::FRT brg1Δ::LEU2 Calb/brg1Δ::HYG* | This study |
| CEM236 | *C. tropicalis* | MYA3404 | *leu2Δ::FRT/leu2Δ::FRT brg1Δ::LEU2 Calb/brg1Δ::HYG* | This study |
| CEM237 | *C. tropicalis* | MYA3404 | *leu2Δ::FRT/leu2Δ::FRT ndt80Δ::LEU2 Calb/ndt80Δ::HYG* | This study |
| CEM238 | *C. tropicalis* | MYA3404 | *leu2Δ::FRT/leu2Δ::FRT ndt80Δ::LEU2 Calb/ndt80Δ::HYG* | This study |
| CEM241 | *C. tropicalis* | MYA3404 | *leu2Δ::FRT/leu2Δ::FRT tec1Δ::LEU2 Calb/tec1Δ::HYG* | This study |
| CEM242 | *C. tropicalis* | MYA3404 | *leu2Δ::FRT/leu2Δ::FRT tec1Δ::LEU2 Calb/tec1Δ::HYG* | This study |
| CEM279 | *C. tropicalis* | MYA3404 | *leu2Δ::FRT/leu2Δ::FRT efg1Δ::LEU2 Calb/efg1::HYG* | This study |
| CEM280 | *C. tropicalis* | MYA3404 | *leu2Δ::FRT/leu2Δ::FRT efg1Δ::LEU2 Calb/efg1::HYG* | This study |
| CEM283 | *C. tropicalis* | MYA3404 | *leu2Δ::FRT/leu2Δ::FRT efg1Δ::LEU2 Calb/EFG1-13Myc-FRT* | This study |
| CEM284 | *C. tropicalis* | MYA3404 | *leu2Δ::FRT/leu2Δ::FRT efg1Δ::LEU2 Calb/EFG1-13Myc-FRT* | This study |
| CEM323 | *C. tropicalis* | MYA3404 | *leu2Δ::FRT/leu2Δ::FRT rob1Δ full ORF::LEU2 Calb/rob1Δ::HYG* | This study |
| CEM324 | *C. tropicalis* | MYA3404 | *leu2Δ::FRT/leu2Δ::FRT rob1Δ full ORF::LEU2 Calb/rob1Δ::HYG* | This study |
| CEM355 | *C. tropicalis* | MYA3404 | *leu2Δ::Calb LEU2/leu2Δ::FRT BCR1/BCR1-13Myc-FRT* | This study |
| CEM356 | *C. tropicalis* | MYA3404 | *leu2Δ::Calb LEU2/leu2Δ::FRT BRG1/BRG1-13Myc-FRT* | This study |
| CEM357 | *C. tropicalis* | MYA3404 | *leu2Δ::Calb LEU2/leu2Δ::FRT EFG1/EFG1-13Myc-FRT* | This study |
| CEM358 | *C. tropicalis* | MYA3404 | *leu2Δ::Calb LEU2/leu2Δ::FRT NDT80/NDT80-13Myc-FRT* | This study |
| CEM359 | *C. tropicalis* | MYA3404 | *leu2Δ::Calb LEU2/leu2Δ::FRT ROB1/ROB1-13Myc-FRT* | This study |
| CEM360 | *C. tropicalis* | MYA3404 | *leu2Δ::Calb LEU2/leu2Δ::FRT TEC1/TEC1-13Myc-FRT* | This study |
| CEM367 | *C. tropicalis* | MYA3404 | *leu2Δ::FRT/leu2Δ::FRT flo8Δ::LEU2 Calb/flo8Δ::HYG* | This study |
| CEM368 | *C. tropicalis* | MYA3404 | *leu2Δ::FRT/leu2Δ::FRT flo8Δ::LEU2 Calb/flo8Δ::HYG* | This study |
| CEM429 | *C. tropicalis* | MYA3404 | *leu2Δ::Calb LEU2/leu2Δ::FRT FLO8/FLO8-13Myc-FRT* | This study |
| NRRL Y-11826 | *C. lusitaniae* | NRRL Y-11826 | | Phaff collection |
| NRRL Y-7426 | *D. hansenii* | NRRL Y-7426 | | Phaff collection |
| NRRL YB-4239 | *L. elongisporus* | NRRL YB-4239 | | Phaff collection |
| NRRL Y-17104 | *P. stipitis* | NRRL Y-17104 | | Nocedal I. *et al*, 2017 |
| NRRL Y-324 | *M. guilliermondii* | NRRL Y-324 | | NRRL collection |
| NRRL Y-1498 | *C. tenuis* | NRRL Y-1498 | | NRRL collection |
| NRRL Y-27907 | *S. passalidarum* | NRRL Y-27907 | | NRRL collection |
| L7554/TBR1 | *S. cerevisiae* | Sigma1278b | *ura3-52 his3::hisG leu2::hisG* | Gerald R. Fink |
| CBS138 | *C. glabrata* | CBS138 | | Sheena Singh-Babak |
